# Molecular characterization of unique multi-domain harbouring fungal rhodopsin for establishing their novel opto-synthetic biological usages

**DOI:** 10.1101/2025.06.26.661701

**Authors:** Alka Kumari, Abhishek Kumar, Komal Sharma, Swaroop Ranjan Pati, Shilpa Mohanty, Suneel Kateriya

## Abstract

Organisms employ light as an external stimulus for regulating cellular functions. The light-sensitive photoreceptors detect light at varying wavelengths, activating signaling cascades and triggering a range of physiological responses. Rhodopsin is a transmembrane heptahelical protein that functions as an ion channel, or a pump, and sensory receptor, respectively. It consists of a light-sensing chromophore, a retinal that upon absorbing light, initiates a series of signaling pathways of sensory perception, growth and survival. Modular rhodopsin (Different from Rhodopsin-Cyclase Module) has been reported in lower eukaryotes, its identification, characterisation and functional significance in the Fungal Kingdom largely unknown. Here, we report the identification of novel modular rhodopsins in fungi, which highlights their potential usages towards the unexplored opto-biotechnological applications (e.g., biomanufacturing of terpenoids, cytoskeleton regulation, DNA metabolism, light-controlled acetyltransferase, etc.) simply by illumination. Furthermore, identification of novel modular rhodopsins augments the expansion of the new optogenetic tools for a wide range of relevant applications. The structural and homology analysis of these identified domains sheds light on their evolutionary lineage and relatedness with the well-characterised bacteriorhodopsin, sensory and channelrhodopsin. The interactome analysis effector domain coupled with the microbial rhodopsin (Rh) reveals RPEL-mediated gene expression and metabolite regulation, which further modulates the retinol synthesis pathway. The role of the fungal Rh-RPEL effector domain in modulating the terpenoid and sphingolipid metabolism in response to light was successfully elucidated via protein-protein interaction and Biosynthesis Gene Cluster (BGC) analysis. This highlights the potential of these novel opto-synthetic biological usages that can induce the light-dependent production of commercially relevant fungal bioactive(s).

## 1. Introduction

The sunlight serves as a vital source of energy for the ecosystem, mediating photosynthesis and supporting the survival of non-photosynthetic heterotrophic organisms. Animals do not display direct interaction with light; instead, they utilize light as an environmental cue. The light-sensors in these organisms are the photoreceptors utilizing chromophores as the light-sensing unit. In mammals, light sensing has been widely studied in the eye where rods and cones opsins absorb light at different wavelengths (Rao and Xue, 2024). Apart from mammals, light is a significant environmental signal for several organisms including bacteria, algae, and fungi. Light influences diverse processes such as development and energy storage. The identification of the channelrhodopsin (ChR) for the first time, in the green microalgae, *Chlamydomonas reinhardtii,* served as an accelerator for the study of photoreceptors (Emiliani et al., 2022).

Rhodopsin is a heptahelical transmembrane protein that is ubiquitously found in all domains of living organisms. It consists of opsins, apoproteins and the vitamin A-derived chromophore, retinal that absorb light, triggering a range of signaling processes for adaptation and survival. Rhodopsin undergoes activation by photoisomerization of retinal chromophore that is covalently linked to the lysine residue via a protonated Schiff base. The opsin (apoprotein) shows variability in the acceptance of retinal isomers, leading to the formation of pigment analogs. The rhodopsins are divided into two categories-microbial or type-I and animal or type-II. However, both the type of rhodopsins share a common structure involving seven transmembrane α-helices. The microbial rhodopsins employ retinal in the all-*trans*/13-*cis* configurations, whereas in contrast animal rhodopsins adopt a 11-*cis*/all-*trans* arrangement (Liu and Colmenares, 2003; Ernst et al., 2014). The type-II rhodopsins belong to the group of G-protein coupled receptors (GPCRs) that convert light energy into electrophysiological signals for vision. However, the type-I rhodopsins, such as bacteriorhodopsin (BR from *Halobacterium salinarum*), mediate the conversion of photons into electrochemical energy by transporting H^+^ ions in a light-dependent process. In addition, these microbial rhodopsins serve as channels, ion pumps and signaling enzymes/opsins, all of which are regulated by light (Nagata and Inoue, 2021; Kulbay et al., 2024).

Similar light-sensing mechanisms have evolved in multiple kingdoms such as archaea, fungi, as well as humans. The presence of light-sensors in the saprophytic fungi is indeed an intriguing feature. Filamentous fungi are sessile, where the spores serve as primary propagation units into the ecosystem. However, spore formation is initiated when a fungus grows at the air-water interface, where light serves as a crucial environmental signal for asexual conidia formation, pigmentation, and circadian clock regulation. Hence, fungi have evolved to detect a range of wavelengths of light, starting from ultraviolet (UV) to far-red. The retinal-binding proteins, opsins possess seven transmembrane helices that absorb light for mediating signaling and energy conservation. Although eyes are primarily absent in fungi, rhodopsins have been identified in *Neurospora crassa* (NR-NOP-1), *Leptosphaeria maculans* and *Allomyces reticulatus* (Furutani et al., 2004; Waschuk et al., 2005; Purschwitz et al., 2006).

The presence of retinal in tissues makes rhodopsin a suitable candidate for optogenetic modulation to control biological function by light. Light serves as an excellent agent for optogenetic-based photodynamic therapy as it displays low toxicity, enhanced spacio-temporal control and regulate drug activation by tempering duration, intensity and wavelength of the light (Vickerman et al., 2021). Recent metagenomic research has shown that various organisms, including fungi and flagellates, possess numerous rhodopsin-like genes. The research team identified rhodopsin (Rh) featuring a phosphodiesterase (PDE) module (Rh-PDE) capable of hydrolysing intracellular second messengers cAMP and cGMP within the flagellate genome, marking the first report of a light-activated phosphodiesterase globally (Yoshida et al., 2017). Furthermore, fungi that produce zoospores, such as *Blastocladiella emersonii* and *Rhizoclosmatium globosum*, contain rhodopsin (Rh) equipped with a guanylate cyclase (GC) module (Rh-GC), which facilitates light-triggered signaling by generating cGMP in response to light. It is believed that this signaling plays a role in the formation of zoospores induced by light (Scheib et al., 2015). Therefore, the modular rhodopsins have evolved in the microorganisms by utilising light and performing a wide array of functions.

The discovery and identification of novel modular rhodopsins paves way for designing new optogenetic tools that can be harnessed for biotechnological applications. With this objective in view, in this study, we examined the fungal genome database, MycoCosm and discovered multiple new modular rhodopsins, i.e. Rh coupled with different effector domains (Rh-RPEL, Rh-MCM, and Rh-NADB-Rossmann, Rh-GC-Carn_acyltrans). Each of these effector domains has its unique function, independent of each other but somehow regulated by light.

For the first time, the novel modular domains in fungi like RPEL repeats, NADB-Rossmann domain and MCM binding domain have been reported that are coupled with rhodopsin. The identification of these novel effector domains in the kingdom fungi highlights an immense optogenetic potential. Not only in green lineage, but also in microorganisms such as fungi, light can be used to modulate the structural biology, energy metabolism and cellular activities such as transcription. Further, the homology and structural analysis of these newly screened modular rhodopsin gives suggestive data regarding its course of evolution. This unexplored branch of optogenetics sheds insight into the regulatory role of light in the microbial world.

## 2. Materials and Methods

### 2.1 Rhodopsin sequence identification and analysis of the novel modular rhodopsins from fungal genome

The reference protein (bacteriorhodopsin) containing the rhodopsin domain was obtained from the National Center for Biotechnology Information (NCBI). The sequences were utilized as queries for BLASTp and tBLASTn searches in the NCBI non-redundant protein sequence database and core nucleotide database, respectively, as well as in the JGI MycoCosm database (https://mycocosm.jgi.doe.gov) as of 31^st^ May 2025. We have selected the sequence for analysis that exhibits a significant E-value. Additional searches were conducted against the Protein Data Bank (PDB) and the transmembrane protein database (PDB_TM) (Tusnády et al., 2005). The NCBI conserved domain search was utilized to identify conserved motifs by comparing the acquired sequences with the NCBI curated database (Wang et al., 2023). The sequences of rhodopsin were aligned utilizing Clustal Omega with the standard settings (Madeira et al., 2024). The primary aim was to identify rhodopsin domains featuring the conserved seven transmembrane domains. These rhodopsins were subsequently examined for functionally conserved critical amino acid residues and retinal binding motifs. These motifs were used to identify and classify these novel modular rhodopsins. The location of an amino acid is indicated with respect to BR, if stated differently in the manuscript and presented in table form **(Supplementary Table 1)**. The global alignment affects the homology of the conserved amino acid residues in the rhodopsin TM7 region. We examined the existence of crucial conserved residues through the BioEdit software (Hall, 2011). To study the potential existence of a retinal chromophore synthesis pathway, the enzymes involved in carotenoid biosynthesis present in *B. emersonii* genome (Galindo-Solís et al., 2022) were utilized. The protein sequences from this collection were subjected to the BLAST search (Altschup et al., 1990) (BLASTp) against the NCBI and Mycosom databases.

### 2.2 Phylogenetic pattern of the novel fungal modular rhodopsins (different from rhodopsin-cyclase module)

The modular rhodopsins retrieved from JGI and NCBI were aligned with different well-characterized microbial rhodopsins associated with different families. The diverse microbial rhodopsins include bacteriorhodopsin (BR), channelrhodopsin (ChR2), and anabaena sensory rhodopsin (ASR). The newly retrieved sequences were aligned by the Clustal ω program of BioEdit against the above-mentioned well-characterized protein sequences. The output of analysis by BioEdit was utilized for the phylogenetic analysis through MEGA X software by the Neighbour-joining (NJ) method using thousand bootstrap replicates (Kumar et al., 2018). Topology was visualized by tree view and NJ plot (Perrière & Gouy, 1996).

### 2.3 Biocuration of Protein-Protein Interactome (PPI) of newly identified multi-domain containing fungal rhodopsins

The protein-protein interactions of effector domain(s) in association with rhodopsins, i.e., RPEL, MCM, and NADB, were predicted by the String database (Mering et al., 2003). The predicted protein-protein interactome obtained as an output, that was subsequently used to generate the network via Cytoscape 3.7.2 (Shannon et al., 2003; Saito et al., 2012). The genome sequence of the ascomycete, *Aureobasidium pullulans* was downloaded from NCBI database and was employed as a template for the identification of Biosynthetic Gene Cluster (BGCs) across its entire genome via antiSMASH (Blin et al., 2016) fungal version. The genome sequence was uploaded in the FASTA format and annotation were in the *.gff format. The default parameters were used for the detection of biosynthetic genes and other supporting proteins (enzymes, transporters and cluster borders). The regulatory and functional interactions of BGC genes were analyzed by protein-protein interaction (PPI) network in the STRING database. Each biosynthetic gene was individually used as a query to predict interaction based on the literature studies, co-expression, neighbourhood, and text mining. The predicted PPI network was exported to Cytoscape v 3.10.3 (Saito et al., 2012) for further analysis. The genes were subjected to functional enrichment and were grouped accordingly.

To explore the light-based modulation of BGCs, the rhodopsin RPEL, light sensitive modular photoreceptor was introduced in the interaction network. The objective of such inclusion was to monitor the possible metabolic flux modulation by light. The metabolic pathways liked to terpenoid, carotenoid and sphingolipid biosynthesis were prioritized for the opto-synthetic modulation of biosynthetic compounds.

### 2.4 Structure-function determinants and homology modelling of the fungal rhodopsin domain: An *Ab initio* structure prediction, molecular docking and structural homology

Ab initio three-dimensional (3D) structure prediction was accomplished with help of Alphafold2 Colab (David et al., 2022). Structure preparation of the predicted structure was achieved prior to molecular docking. Hydrogen atoms were added and Gasteiger charges were computed using dockprep program of UCSF chimera. The 3D structure of all-trans-retinal was retrieved from PubChem database (CID: 638015) (Kim et al., 2025). The ligand structured was optimised by structure minimization algorithm in UCSF chimera. The coordinates of retinal binding pocket were as per the reference bacteriorhodopsin (PDB ID: 1XIO) close to the lysine residue of 7^th^ transmembrane helix for the formation of Schiff base bond. Possible interaction of all-*trans* retinal was predicted by molecular docking with help of UCSF chimera and AutoDock Vina (Pettersen et al., 2004; Trott & Olson, 2010). The resulting protein-ligand interactions were ranked based on the binding affinity scores. The top ranked poses were selected for interaction analysis via PyMOL (Schrödinger & DeLano, 2020).

Protein-ligand interaction profiler (PLIP) was used to study the possible interaction of retinal molecule and rhodopsin inside retinal binding pocket (Salentin et al., 2015). The Swiss-model was used for structural homology analysis. The structure was compared on Swiss-model platform and comparative overlapping was achieved with help of UCSF chimera (Bienert et al., 2017; Pettersen et al., 2004; Waterhouse et al., 2018).

## 3. Results and Discussion

### 3.1 *In silico* mining of fungal genome reveals four different modular rhodopsins

Mining the fungal genome database uncovered four distinct types of MTRs (microbial-type rhodopsins) occupying different habitats. Continuous advancements in understanding the photobehavioral response of *Chlamydomonas* directed the initial identification of modular rhodopsins, but subsequent discoveries in other organisms have been limited. Recent metagenomic studies have revealed that numerous organisms, such as fungi and flagellates, harbour many rhodopsin-like genes. In this way, microorganisms exploit light through rhodopsins that fulfil various roles. The MycoCosm (fungal) genome database was thereby explored. We discovered new microbial modular bacteriorhodopsin **(Figure 1A and Table 1)**, modular channelrhodopsin-like sequence **(Figure 1A-B and Supplementary Table 1)**, and twin modular rhodopsin **(Supplementary Figure 1)** within different fungi. We evaluated the key characteristics differentiating MTRs from the other seven transmembrane protein families. In the twin rhodopsin, it was observed that there were variations in the number of transmembrane helices, as the number varied between 5 and 6. However, they were flanked by multiple transmembrane helices, whose function remains ambiguous **(Supplementary Figure 1)**. As literature studies have shown that GC as a modular domain has been widely studied, therefore, this work targets the unravelling of the other novel effector domains.

**Figure 1:**
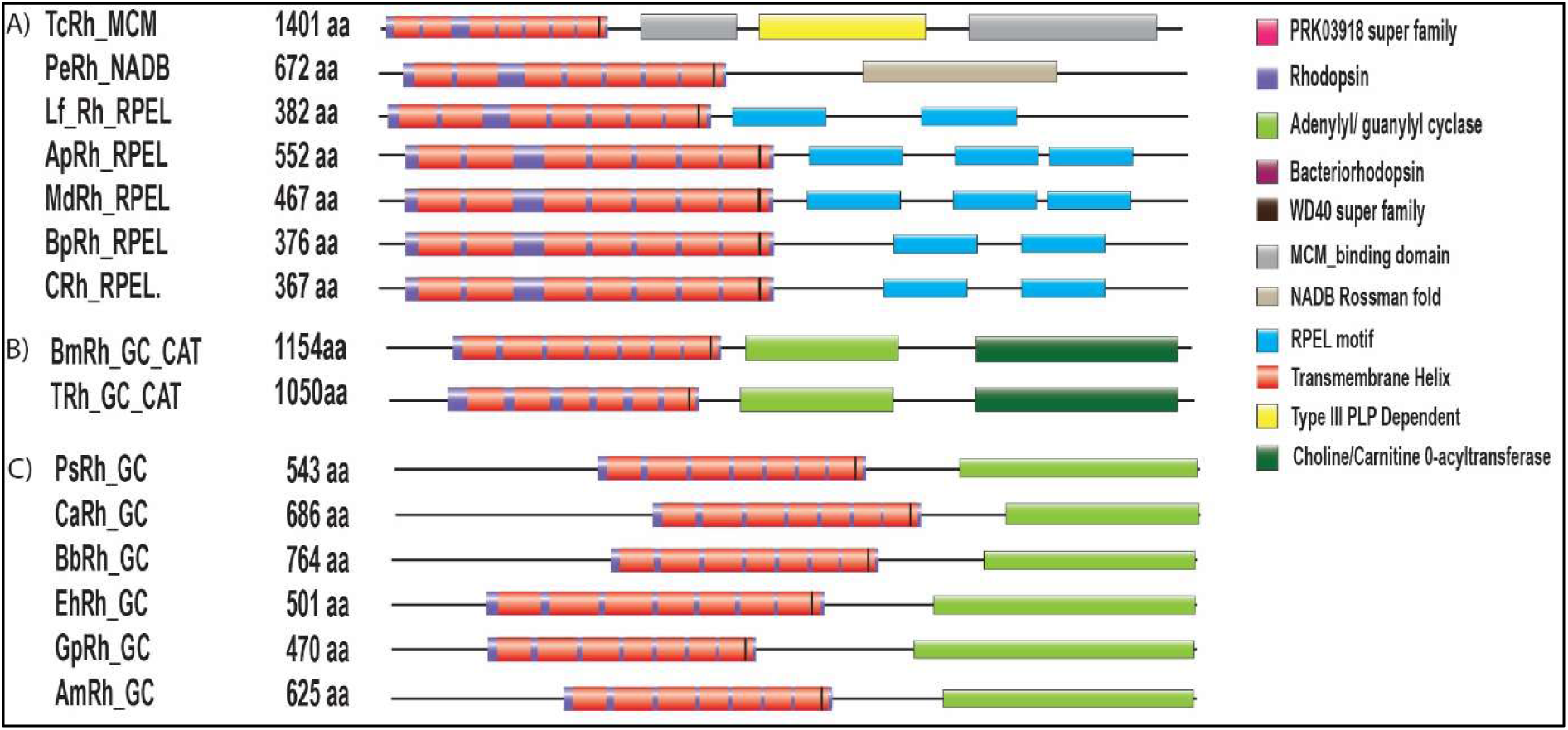
The schematic illustration depicts rhodopsin with modular domains, the blackline denotes the full-length protein, and domains are illustrated by geometric structures. **(A)** Domain organization of modular Bacteriorhodopsins (BRs), coupled with MCM (Minichromosome Maintenance) in *Testicularia cyperi* (TcRh), (NAD(P)-Binding Rossmann Fold) in *Pseudographis elatina* (PeRh) and RPEL repeats in *Lenthithecium fluviatile* (LfRh), *Aureobasidium pullulans* (ApRh), *Myriangium duriaei* (MdRh), *Baudinia Panamericana* (BpRh), *Capnodiales* sp. (CRh) **(B)** Domain organisation of modular Channelrhodopsin-like sequence (ChRs) coupled with CRAT (Carnitine O-acetyltransferase) in *Boothiomyces macroporosus* (BmRh), *Terramyces* sp. (TerspRh). **(C)** Domain organisation of modular Channelrhodopsin-like sequence coupled with GC (Guanylate Cyclase) in *Paraphysoderma sedebokerense* (PsRh), *Caternaria anguillulae* (CaRh), *Blastocladiella britannica* (BbRh), *Entophlyctis helioformis* (EhRh), *Globomyces pollinis-pini* (GpRh), *Allomyces macrogynus* (AmRh).

### 3.2 Characterization of identified sequences for functional prediction of light-gated ion channels, sensory and ion pumps based on conserved residues of the microbial rhodopsins

Rhodopsin activation and functions are mostly determined by the position of the absolute amino acid located close to the retinal binding site. Channelrhodopsins and bacteriorhodopsin are types of microbial rhodopsins that utilize retinal as a chromophore and experience light-induced structural changes; however, they vary in their proton pathways and functional results.

#### 3.2.1 Identification of novel fungal modular light-gated ion pump

The search for the modular BR resulted in three modular BRs as depicted in **Figure 1A**. These are Rh-RPEL repeats, Rh-NAD(P)-Binding (NADB) Rossmann Fold and Rh-MCM from different fungi in the ascomycete phylum. The optogenetic applicability of these modular domains (RPEL repeats, NAD(P), and MCM) are summarised in **Supplementary Table 2**. The rhodopsin domains of Rh-RPEL, Rh-NAD(P), and Rh-MCM were aligned with well-characterized BR and ASR (*Anabaena* sensory rhodopsin) taken as the reference sequence. The conserved residues crucial for the photocycle are highlighted in **Figure 2**, and the same have been studied for major functionalities, which are: (1) The photoisomerization of retinal, linked to Lys216 by a Schiff base, initiating a sequence of proton transfers. (2) The Schiff base (SB) transfers its proton to Asp85. (3) Asp96 then donates a proton to restore the Schiff base from the cytoplasmic side (Luecke et al., 2001). Arg 82, Glu194, and Glu204 are involved in facilitating the release of protons towards the extracellular side. Asp212 aids in maintaining the correct electrostatic environment necessary for effective proton transfer. Depending on these amino acid residues, the rhodopsin domains were analysed and the details of the summary is in **Supplementary Table 2**. The boundary of the specific helices was taken according to the bacteriorhodopsin sequence, containing seven hydrophobic alpha-helices with a conserved lysine in the G-helix that forms a retinal binding pocket. The identified BR-like sequences showed the same triad residues corresponding to BR in the C-helix, except TcRh, which has DTE corresponding to BR’s DTD. The ion specificity of a rhodopsin is determined by the third position of the triad residue. We observe valine (PeRh, ApRh), leucine (TcRh), and in other cases, isoleucine at the BR-G116 analogue site, which is typically considered a distinguishing feature of fungal rhodopsins **(Figure 2, Supplementary Table 1)**. The group responsible for proton release (BR: E194, E204, D212) is highly conserved across the mentioned modular rhodopsins; R82 primarily participates in the release of protons to the extracellular side, showing a strong conservation among the reported rhodopsins. These rhodopsins appear to be operational because the lysine that binds retinal is preserved across all of them. In this report, we also discuss an HR-like sequence found in *Cladophialophora carrionii*, which is a significant cause of human chromoblastomycosis. The ClacRh exhibits similarity to BR but includes an NTQ motif, characteristic of halorhodopsin. The initial light-driven inward chloride pump was identified as HR in the year 1977 (Matsuno-Yagi and Mukohata, 1977) and features the TSA motif. NTQ rhodopsins, such as *Fulvimarina* rhodopsin (FR), with the D85 position substituted by asparagine, demonstrate Cl^−^-dependent colour variations, signifying direct interactions near the Schiff base, similar to HR (Inoue et al., 2014). The validation of the amino acids in the retinal-binding pocket has been studied using a mutagenesis approach, where the findings have been summarised in **Supplementary Table 3**.

**Figure 2:**
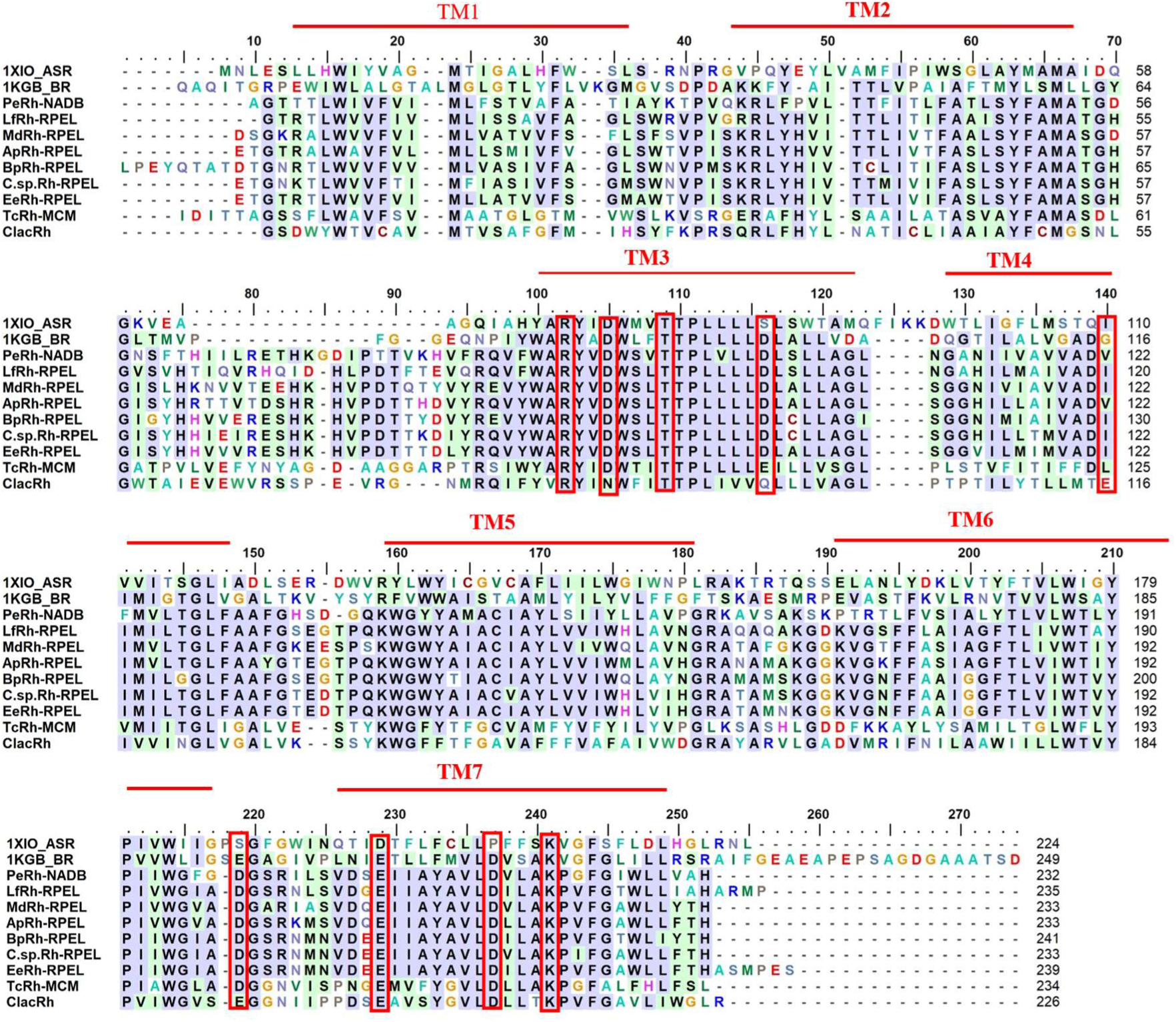
A multiple sequence alignment of microbial rhodopsins from different fungi highlighting the conserved residues. Identical amino acids are marked in blue, indicating a higher level of sequence conservation, whereas similar amino acids are highlighted in lighter green, suggesting functional or structural conservation despite variation in sequence. The alignment covers the seven-transmembrane (7TM) regions. The location of the lysine residue that binds retinal and the spectral tuning residues are indicated by a red square. The alignment has been evaluated using bacteriorhodopsin protein as the template. 1KGB: Bacteriorhodopsin, 1XIO: *Anabaena* sensory rhodopsin.

#### 3.2.2 Identification of novel fungal modular channelrhodopsin-like sequences

A new type of rhodopsin displays a distinctive arrangement of domains with the rhodopsin located at the N-terminus within a single open reading frame, with guanylate cyclase and carnitine o-acyltransferase (Rh-GC-CRAT) at the C-terminus in *Boothiomyces macroporosus* and *Terramyces* sp., (microscopic chytrid fungus). The rhodopsin domain of BmRh, Tersp, is used to determine the functionality. The rhodopsin domains were aligned with the well-studied ChR2, taken as the reference. The key amino acids crucial for the photocycle have been examined for their primary roles, which include (1) the lysine that binds to the retinal, (2) the counter ion or proton acceptor of the retinal Schiff base (RSB), (3) the maintenance of the proton acceptor, and (4) the DC-gate located in helices 3 and 4. The lysine residue located at the 7^th^ helix forming a covalent bond with retinal are conserved **(Figure 3)**. The BmRh possesses the conserved seven transmembrane domains. However, the TerspRh lacks the 7^th^ transmembrane helix but possesses the conserved lysine residue that may impact the formation of the retinal binding pocket. The aspartate residue located at the 156^th^ position in ChR2 provides a proton to the RSB during the re-protonation process, and these interactions are maintained in Tersp-Rh, but in BmRh, they are substituted with Asn(N) weakens cation coordination **(Figure 3)**. Asp156 in ChR2 forms a hydrogen bond with Cys128, creating a DC-gate as a switch for ion movement (Miralles et al., 2003). Cys128 is also retained in the newly identified rhodopsin **(Figure 3)**. Consequently, the functional residues in BmRh and TerspRh that correspond to ChR2 demonstrate substitutions such as E90-N and E97-Y (the pore-gating glutamates), as well as an internal proton donor in BmRh at D156-N and H137-L **(Figure 3)**. The presence of hydrophobic transmembrane domains with aromatic residues may create a positively charged environment that enhances anion selectivity. BmRh-GC-CRAT includes analogs of all critical proton-handling residues found in ChR2, although their specific positioning and interactions might vary. This reinforces the notion that it likely maintains light sensitivity through retinal-bound lysine. While it might not function as a conventional ion channel, it appears to utilize proton-sensitive residues for conformational signaling. It suggests that BmRh and TerspRh are likely derived from ChR-like ancestors to fulfil photo-signaling functions. Due to its distinctive sequence, further experimental investigation is warranted and could lead to engineering efforts aimed at improving its characteristics. As such, newly identified channelrhodopsin-like sequence hold promise as optogenetic instruments for controlling novel biological pathways.

**Figure 3:**
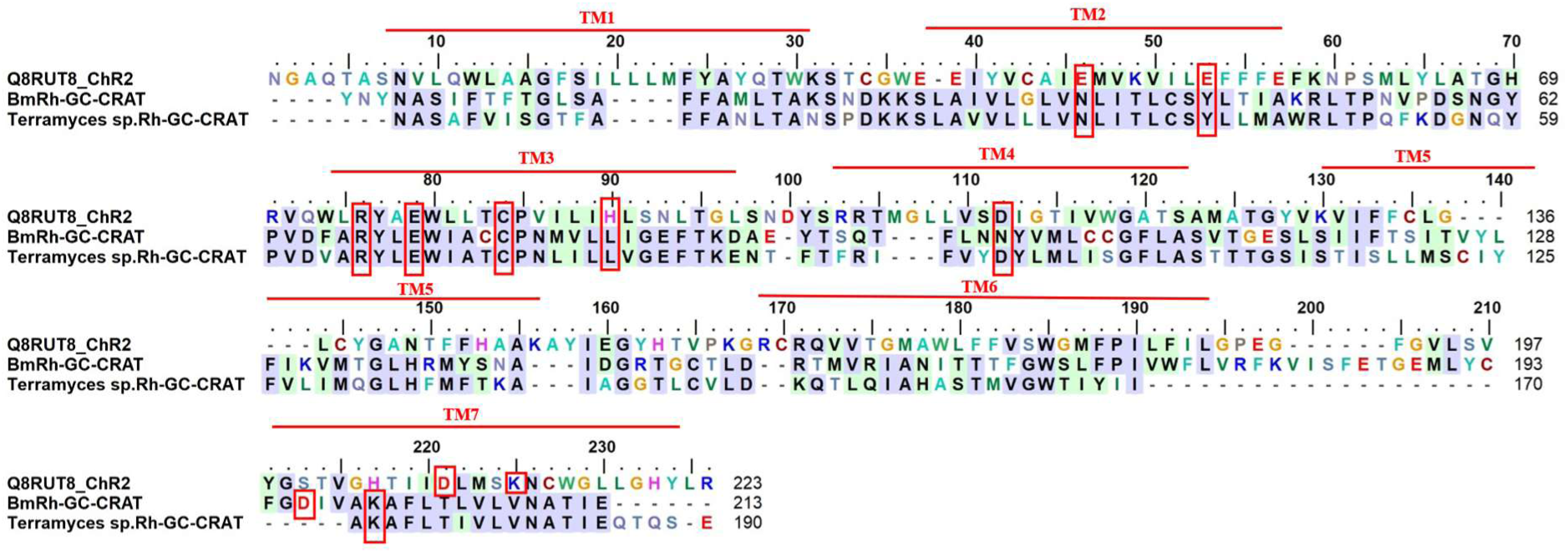
A comparative study of the novel channelrhodopsins and the identification of key amino acid residues: BmRh, TerspRh were aligned with ChR2. Transmembrane (TM1-TM7) are represented by a red bar and numbered accordingly. The retinal-binding lysine, the proton acceptor/donor, the cysteine involved in hydrogen bonding with the proton donor (DC pair), and the arginine crucial for the primary proton translocation are indicated by a red square.

### 3.3 Comparative structural analysis of newly identified fungal rhodopsin

The three-dimensional structure of modular rhodopsin(s) was aligned with the well characterised rhodopsin’s across the biological systems. Upon homology and phylogenetic analysis, sensory rhodopsin (SR1) of *Euglena* (PDB ID: 1XIO) was found to be evolutionary related to the newly found modular rhodopsin. Hence the SR1 protein structure was used as reference for the docking and structural comparison of the newly found rhodopsin. The retinal binding pocket conserved, and affinity score came out to be -9.4 (Docking was repeated thrice, average). The retinal binding pocket was analysed with PLIP and retinal schiff base formation was found along with hydrophobic interaction of other residues forming the retinal binding pocket. Upon structural homology analysis of ApRh utilising Swiss homology model, LmRh was screened as the closest related rhodopsin structurally having light regulated proton pump function. *Aureobasidium pullulans* has already been reported to have light gated proton pump rhodopsin (Panzer et al., 2021). The function of ECL loop has not characterised yet but it has been suggested that it might be helping in stability of rhodopsin in fungal membrane. In this paper we have compared the ECL loop of ApRh with β2 Adrenoreceptor (PDB ID: 2RH1) and corticotropin (PDB ID: 4K5Y) of GPCR family. The variation in loop would be subjected to the specialised function of the rhodopsin and its effector domains **(Figure 4)**.

**Figure 4:**
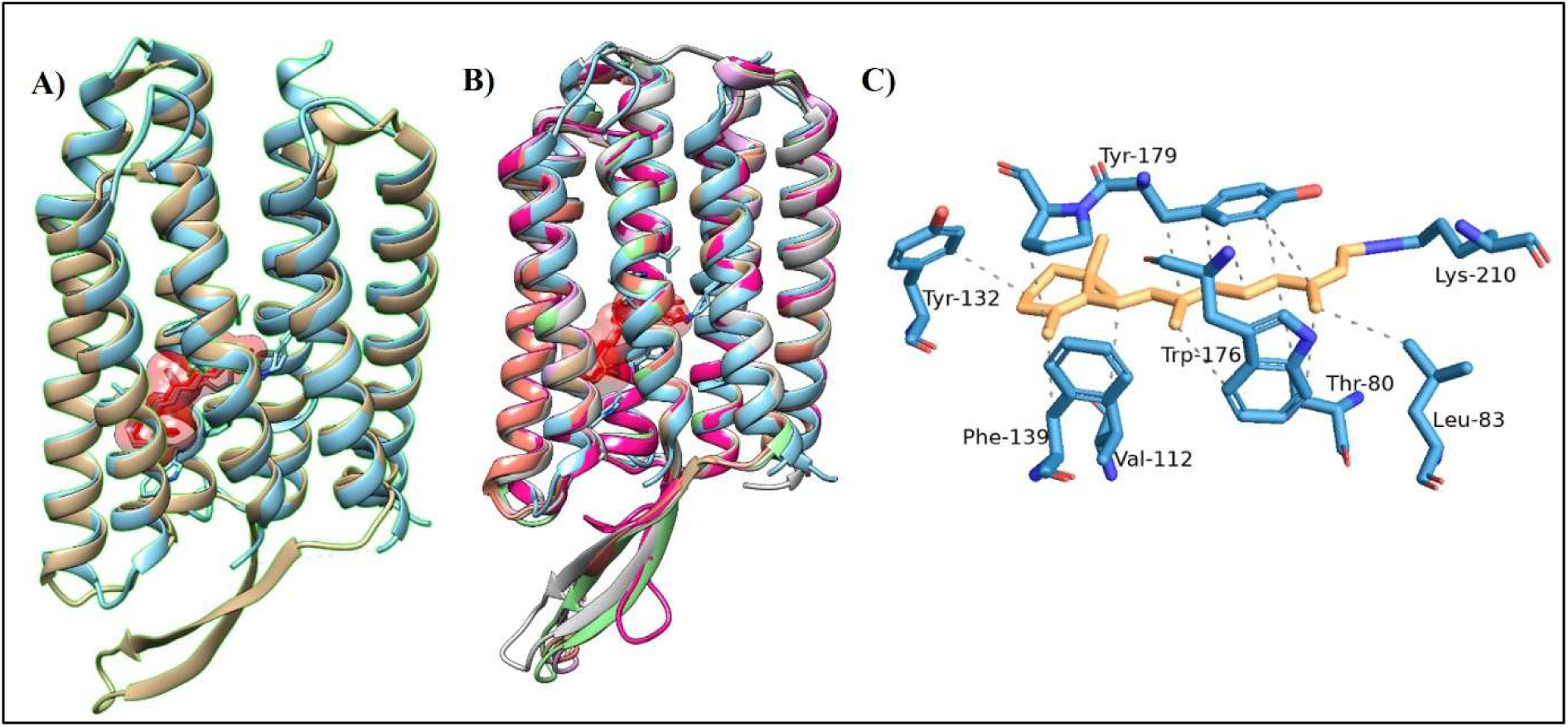
Structural analysis of modular rhodopsin with well characterised bacteriorhodopsin and light gated proton pump. **A)** ApRh (mustard colour) was docked with all-*trans* retinal (red colour) was super imposed with the sensory rhodopsin of *Euglena* (cyan colour). **B)** All-*trans* retinal (orange colour) forming Schiff base bond with lysine residue (blue colour) of rhodopsin of sensory rhodopsin. **C)** ApRh (mustard colour), along with other modular rhodopsins, were superimposed over SR1(PDB ID: 1XIO) and the light-gated proton pump of *Leptosphaeria maculans*.

### 3.4 Modifications in the retinal-binding pockets of modular fungal rhodopsins resulted in a wider range of spectral tuning

Fungal rhodopsins, similar to other microorganisms, exhibit an impressive variety in their spectral characteristics, featuring absorption peaks that span from visible light to near-infrared wavelengths. This characteristic feature is crucial for adapting to different light conditions and carrying out particular functions such as ion transport and zoospore phototaxis. This adjustment occurs through a combination of interactions involving chromophores (retinal) and proteins (rhodopsin). The maximum absorption wavelength (λmax) of microbial rhodopsins is defined by the energy difference between the ground (S0) state and the excited (S1) state of the retinal chromophore, which is covalently attached to the protein through a Schiff base bond. The amino acid residues, Thr89, Ala215, Gly122, Leu93, Asp85, Asp212, Met118, and Trp86 are essential for forming the retinal-binding pocket in bacteriorhodopsin (BR) that impacts its spectral and functional characteristics. Alterations or variations in these residues can result in shifts in absorption maxima, modified photocycles, and changes in the stability or functionality of the protein. A modification in a single nonpolar or polar amino acid at the 105^th^ position in proteorhodopsin can lead to either green-absorbing or blue-absorbing forms of proteorhodopsin, causing a 20 nm shift toward the blue or green spectrum. Mutations like L93A or L93T reduce the photocycle speed by around 100 times relative to the wild-type BR (Kralj et al., 2008). The variance between Ala215 and Thr204 in BR and SRII respectively from *Natronomonas pharaonis* in TM7 accounts for a portion of the 70 nm difference in absorption wavelength between BR and SRII (9 nm of that difference) (Shimono et al., 2001). Some fungal rhodopsins function as heterodimers, consisting of one subunit that captures blue/green light and another subunit (NeoR) that absorbs far-red or near-infrared light. Fungal rhodopsin guanylyl cyclase is a photosensitive enzyme-rhodopsin fusion protein that is vital for zoospore phototaxis in certain fungi like *Blastocladiella emersonii*. Two mutations were introduced with blue-shift (E254D, λmax = 390 nm; D380N, λmax = 506 nm) along with red-shift (D380E, λmax = 533 nm) absorption maxima compared to the wild-type protein (λmax = 527 nm) (Trieu et al., 2017). The replacement of few key residues in novel fungal rhodopsin domain suggests wider spectral tuning of these rhodopsins upon functional expression **(Supplementary Table 3)**.

### 3.5 Evolutionary perspectives of the retinal biosynthesis pathway in relevant fungal genomes

A retinal chromophore cofactor is necessary for rhodopsin function. The molecular processes that underlie the carotenoid proteins’ and light’s control are presently being studied. All photosynthetic organisms, from cyanobacteria to higher plants, synthesize carotenoids, but a variety of heterotrophic microbes, such as fungi and non-photosynthetic bacteria, serve as their producers. Fungal IPP is formed through the mevalonate pathway (Yang et al., 2021). In the initial stages of biosynthesis, prenyl transferases sequentially add IPP (isoprene, C5) units to generate geranyl pyrophosphate (GPP, C10), farnesyl pyrophosphate (FPP, C15), and geranylgeranyl pyrophosphate (GGPP, C20). Retinal, the chromophore of rhodopsin, is produced when the carotenoid oxygenase cleaves β-carotene (Avalos et al., 2017). A retinal chromophore co-factor is necessary for rhodopsin function. The carotenoid (β-carotene) biosynthesis pathway enzyme genes required for retinal formation were found in our selected fungus. We found that the genome database of *Aureobasidium pullulans, Lentithecium fluviatile, Myriangium duriaei, and Testicularia cyperi* contained all three of the enzyme-encoding genes involved in this pathway: bifunctional lycopene cyclase/phytoene synthase, phytoene, and carotenoid oxygenase. The predicted protein-protein interactome for the carotenoid biosynthesis pathway in *Aureobasidium pullulans* is depicted in **Figure 5**. The interactome shows the crosstalk of the major carotenoid biosynthesis enzymes including phytoene desaturase, geranylgeranyl pyrophosphate synthase and carotenoid oxygenase. The abbreviations and the corresponding enzymes have been listed in **Supplementary Table 4**. The key limiting enzymes that are crucial for carotenoid biosynthesis pathway in the fungal genome are detailed in **Supplementary Table 5**. The BLASTp analysis of the phytoene dehydrogenase sequence in *Capnodiales sp.* reveals similarities to the ATP-binding cassette transporter CGR1. Due to the limited information available regarding the fungi *Pseudographis elatina*, *Elasticomyces elasticus*, and *Boothiomyces macroporosus*, the prediction of all three enzymes was not successful.

**Figure 5:**
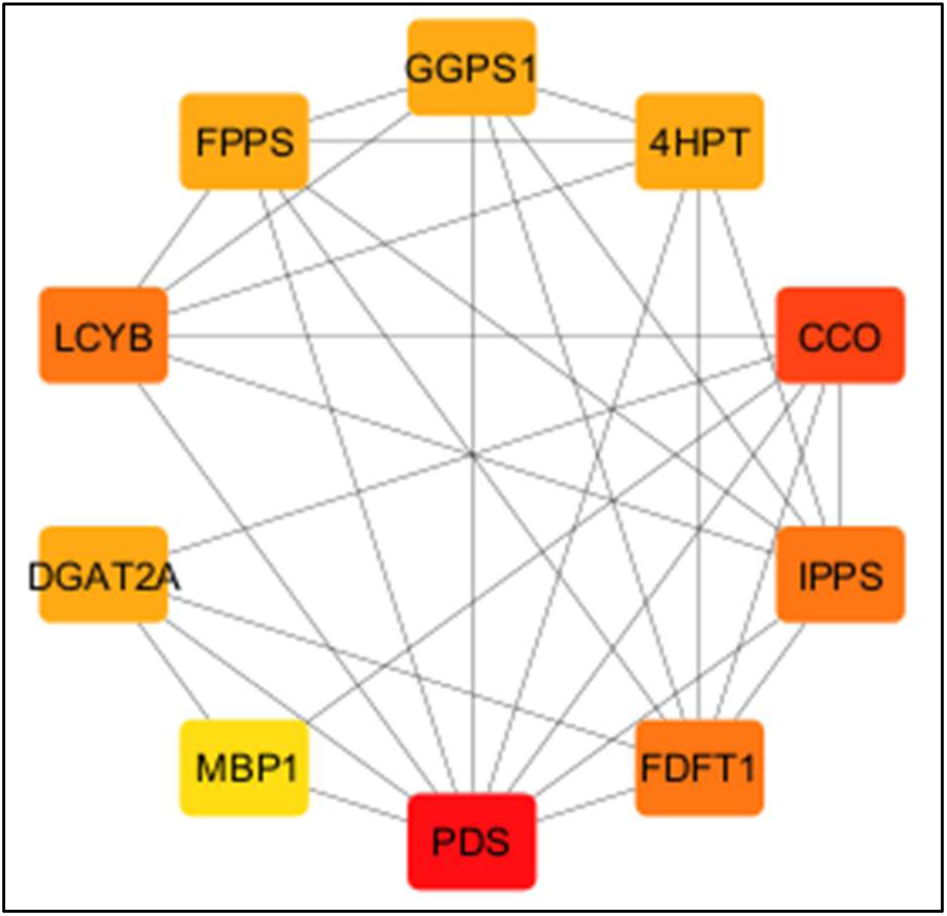
The protein-protein interaction network of the carotenoid synthesis pathway was constructed in *Aureobasidium pullulans* utilizing STRING and visualized through Cytoscape. In this network, nodes symbolize proteins, while edges indicate the interactions between them. Hub proteins were identified by their elevated centrality scores through Cytoscape plugins, CytoHubba, employing algorithms such as degree centrality and shortest path, and were subsequently ranked to determine the most significant proteins in the network. The layout of the network is arranged in a degree-sorted circular fashion. The red nodes represent the proteins with the highest interactions within the PPI, highlighting the centrality of the network, where GGPS1: geranylgeranyl pyrophosphate synthase, 4HPT: 4-hydroxybenzoate polyprenyltransferase, CCO: carotenoid oxygenase, IPPS: Uncharacterized protein, FDFT1: putative farnesyl-diphosphate farnesyltransferase, PDS: phytoene desaturase, MBP1: DNA-binding domain of Mlu1-box binding protein MBP1, DGAT2A: diacylglycerol O-acyltransferase 2A, LCYB: lycopene beta-cyclase and FPPS: ERG20 farnesyl diphosphate synthase **(Supplementary Table 4)**.

### 3.6 Diversification of novel fungal modular rhodopsins uncovers unique evolved lineages

To explore the evolutionary origins of modular rhodopsins, we conducted a phylogenetic analysis to examine their evolutionary lineage, functional variation, and diversification within the microbial world. Classical proton-pumping rhodopsins, such as BR, are distinguished from fusion-type rhodopsins, which indicates their functional divergence. There is a potential new subgroup (RPEL-type) that constitutes a separate cluster. The lower bootstrap value of 44 indicates reduced confidence in those specific branching patterns, which may require additional verification. The phylogenetic tree reveals three distinct clades **(Figure 6).** These are likely rhodopsins featuring RPEL-like motifs and NAD(P) Rossmann folds, possibly functioning differently from standard proton pumps. In the Mid Clade, TcRh-MCM and ClacaRh are grouped with strong bootstrap support of 99. The Lower Clade comprises ChR2 and rhodopsin-GC fusion proteins, forming a separate clade that highlights their divergence from traditional microbial rhodopsins such as BR and ASR. ChR2 and rhodopsin-GC fusion proteins form a distinct clade, indicating divergence from classic microbial rhodopsins like BR and ASR **(Figure 6)**. The adaptation of fungi to their habitats can affect the way they are distributed, specialized, and diversified, influencing their ecological functions and evolutionary paths. *Testicularia cyperi (plant pathogens)* and *Cladophialophora carrionii* (human chromoblastomycosis) are both pathogenic in nature and group in single clades. *Pseudographis elatina* is a drought-resistant, non-lichenized fungus categorized independently of any subclades. Surprisingly, modular rhodopsins grouped themselves without the influence of ASR and BR.

**Figure 6:**
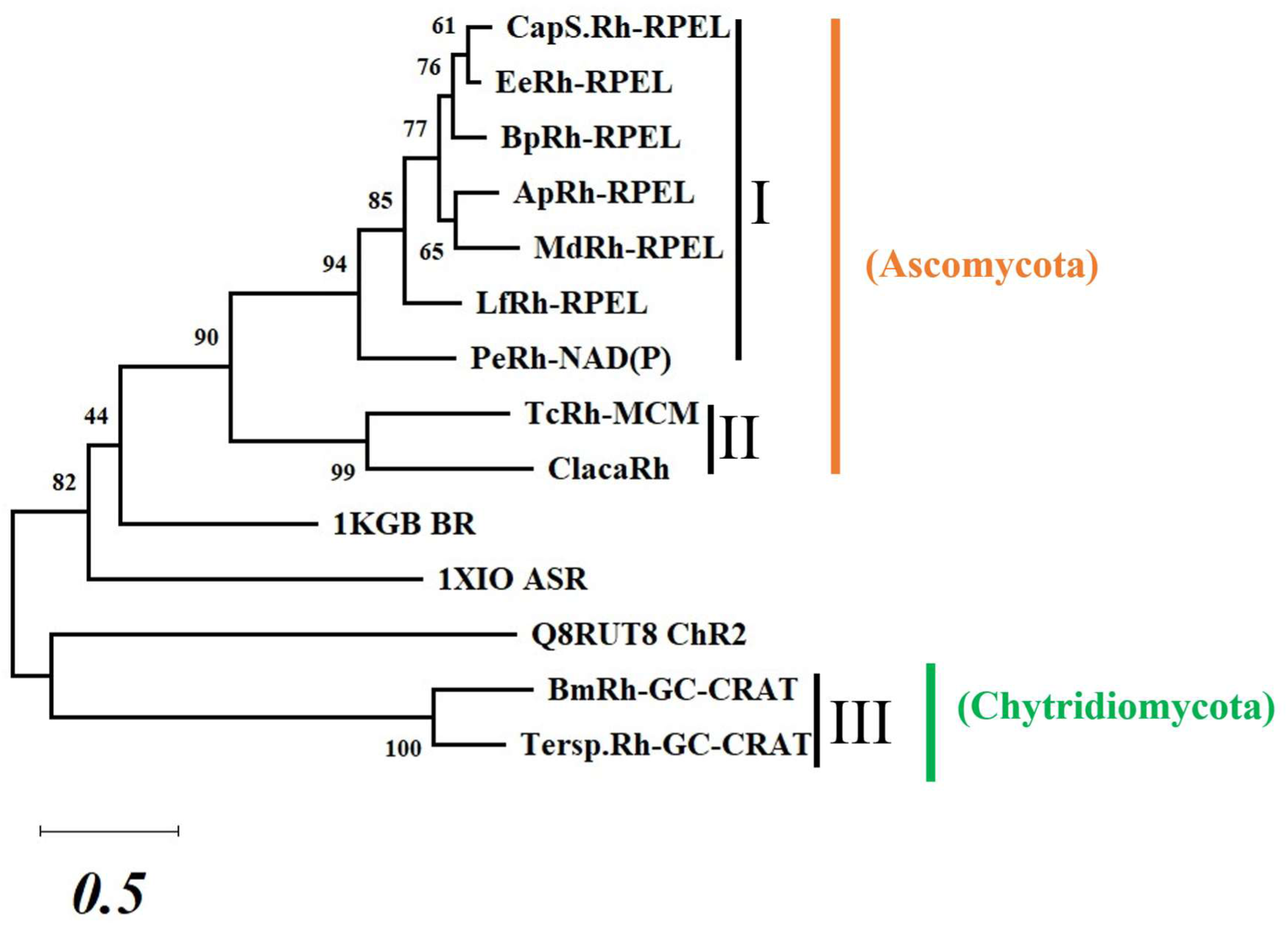
The evolutionary diversification of microbial type rhodopsin was analyzed using the maximum likelihood approach based on the JTT matrix-based model. It incorporates 1000 bootstrap values. The scale bar (0.5) at the bottom represents the branch length, which reflects the degree of sequence divergence. The amino acid sequences that constitute the seven transmembrane helices of rhodopsin were taken into account for pairwise alignment and the creation of a phylogenetic tree. Evolutionary classified into: Ascomycota and Chytridiomycota.

### 3.7 The optogenetic potential of the novel modular fungal rhodopsins

Amongst various effector domains associated with rhodopsin, the highlights were the RPEL repeats, NAD(P) Rossmann fold, and GC-CRAT domains due to their functional relevance, and we evaluated their significance in optogenetics. We performed functional assessments of these domains, and analyzed the protein-protein interaction (PPI) network, and pinpointed their probable interacting members and corresponding pathways **(Figure 7 & 8, Supplementary Figure 2)**. The fundamental biology of MCM proteins in fungi has been extensively investigated in several model organisms, yet there are considerable gaps in understanding regarding early-diverging fungi. For Chytridiomycota, Zygomycota, and Blastocladiomycota, not much is known about the regulation of MCM, which might exhibit distinct regulatory mechanisms. The complete sequence of Rh-MCM was examined using the NCBI conserved domain search program, revealing that it contains three consecutive domains: Rh-(PLP) Pyridoxal 5-phosphate-MCM (Mini-chromosome maintenance replisome factor). For function prediction, we conducted protein-protein interaction (PPI) analysis using the full-length sequence as well as domain-specific analyses. However, we could not find specific PPIs related to MCM biology in the existing literature. To facilitate the potential optogenetic application of Rh-MCM, it is necessary to characterize the MCM biology of these early-diverging phyla.

**Figure 7:**
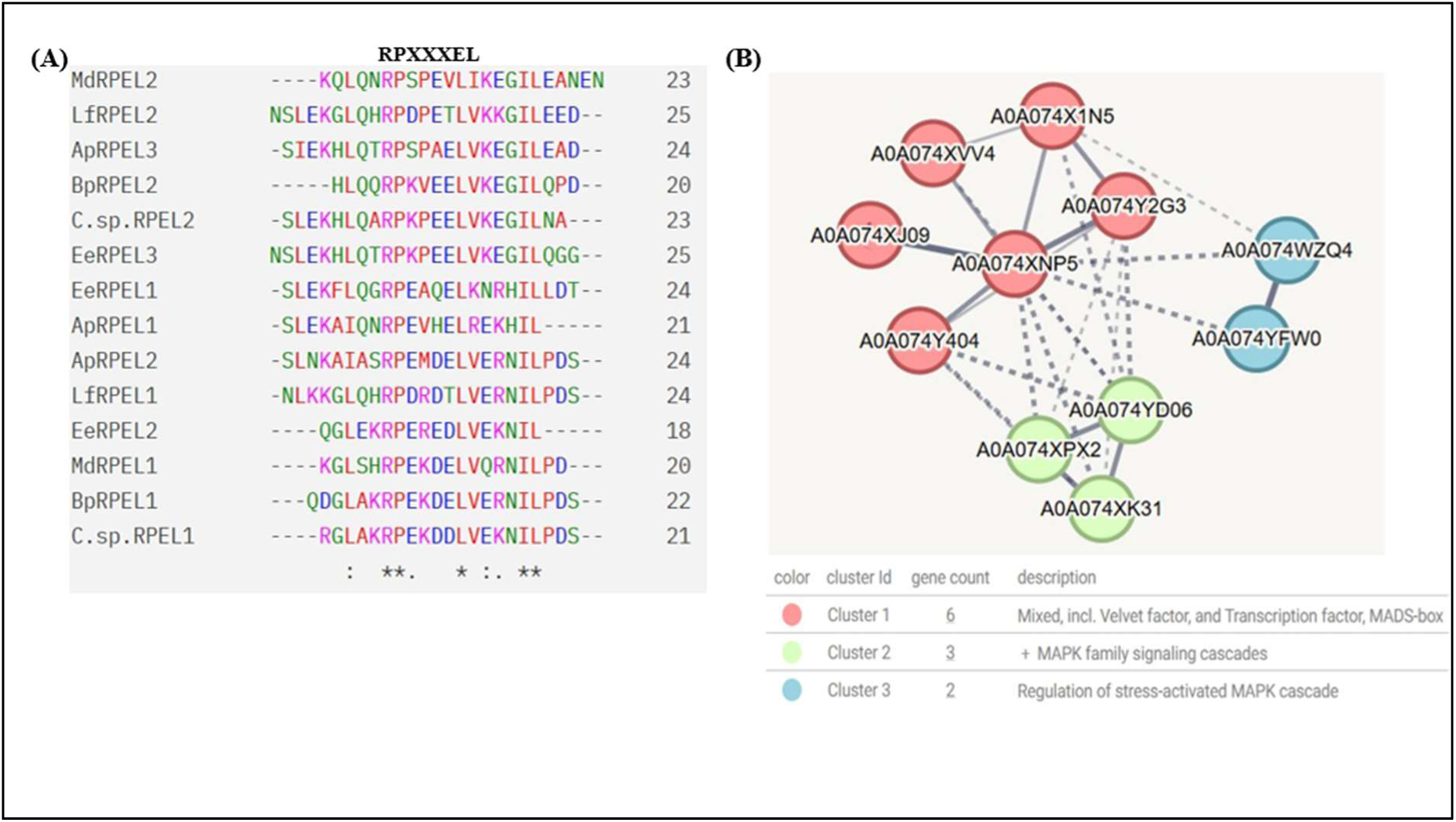
(A) Sequence alignment of individual RPEL motifs from identified modular rhodopsin. Conserved motifs are highlighted by an asterisk. **(B)** Protein-Protein interaction network showing interacting partners of RPEL repeats domains in *Aureobasidium pullulans*. Protein-protein interaction was performed using String version 11 (https://string-db.org/).

#### 3.7.1 Crosstalk of photoreceptors with cytoskeletal elements through RPEL-motif coupled with the rhodopsin domain in Ascomycetes

The characterization of Rh-RPEL repeats is appealing because it may discover an wholly new class of modular rhodopsins. In the NCBI pBLAST search, we found that its presence is confined exclusively to the ascomycetes phylum. G-Protein Coupled Receptors GprM and GprJ in *Aspergillus fumigatus* are essential for regulating the cell wall integrity pathway, producing secondary metabolites, and influencing the organism’s virulence (Filho et al., 2020). Therefore, we hypothesize that a similar biological mechanism will be controlled by the identified modular rhodopsin. RPEL count affects the sensitivity of MRTF to the changes in cellular actin (Watanabe et al., 2012). The proteins BpRh, MdRh, LfRh, CapSp.Rh and EeRh include two RPEL repeat motifs, whereas ApRh features three RPEL repeat motifs **(Figure 1)** according to the conserved domain analysis conducted through NCBI, which needs experimental proof. The sequence homology of RPEL repeats reflects the conserved RPxxxEL motif across the sequences **(Figure 7)**. The protein-protein interaction analysis for the ApRh-RPEL indicated its regulation in the G-actin polymerization biology **(Figure 7)**. The predicted interactome of the RPEL domain was subjected to the CytoHubba analysis **(Supplementary Figure 2)**. It was inferred from the interactome that the RPEL effector domain participates and regulates several metabolic activities within the cell. It interacts with the proteins involved in protein folding and RNA modification within the cell as indicated in the interactome analysis. The RPEL-based interaction with the MAP kinases and chaperones such as Hsp70 regulates gene expression and protein folding kinetics. This further leads to the modification in the composition of the secondary metabolites, as evidenced by the RPEL interaction with proteins involved in sphingolipid metabolism.

#### 3.7.2 Modular nature of rhodopsin–NAD(P) Rossmann fold protein in Pseudographis elatina: A possible connection of light-dependent control of alcohol metabolism

Rhodopsin-NAD(P) Rossmann fold-mediated optobiology" describes a hybrid category of light-sensitive proteins that integrate light detection (through rhodopsin domains) with metabolic or redox functions through NAD(P)-binding Rossmann fold domains. This idea signifies a novel approach in synthetic biology and optogenetics. Our study gives the first report to the scientific group that the rhodopsin-coupled NAD(P) Rossmann fold is naturally present in the fungus *Pseudographis elatina*, which is a non-lichenized fungus. The existing literature does not provide information on the habitat of this fungus. Using structural homology blast, we found that the identified sequence shares homology with short-chain dehydrogenase (7bsx.1). Because there is a lack of genomic or proteomic data in public databases, *Pseudographis elatana* is not featured in the STRING database. Since *Pseudographis elatana* is part of the order Rhytismatales and the class Leotiomycetes (phylum Ascomycota), we searched for similar fungi in STRING. *Botrytis cinerea* possesses well-defined genomes and proteomes in the Leotiomycetes class and is commonly utilized as a model organism for studying fungal interactions. Therefore, enter the blast homolog of *Pseudographis elatana* (NAD(P) into STRING to investigate protein interaction networks within *Botrytis cinerea*, which was selected as the organism of the same class. The PPI analysis shows that the Rh-NAD(P) may regulate alcohol metabolism. Using a BLAST search, we identify homologs of PPI in *Pseudographis elatana*. We can speculate that Rh-NAD(P) could be involved in alcohol metabolism and may influence its non-lichen characteristics **(Figure 8)**.

**Figure 8:**
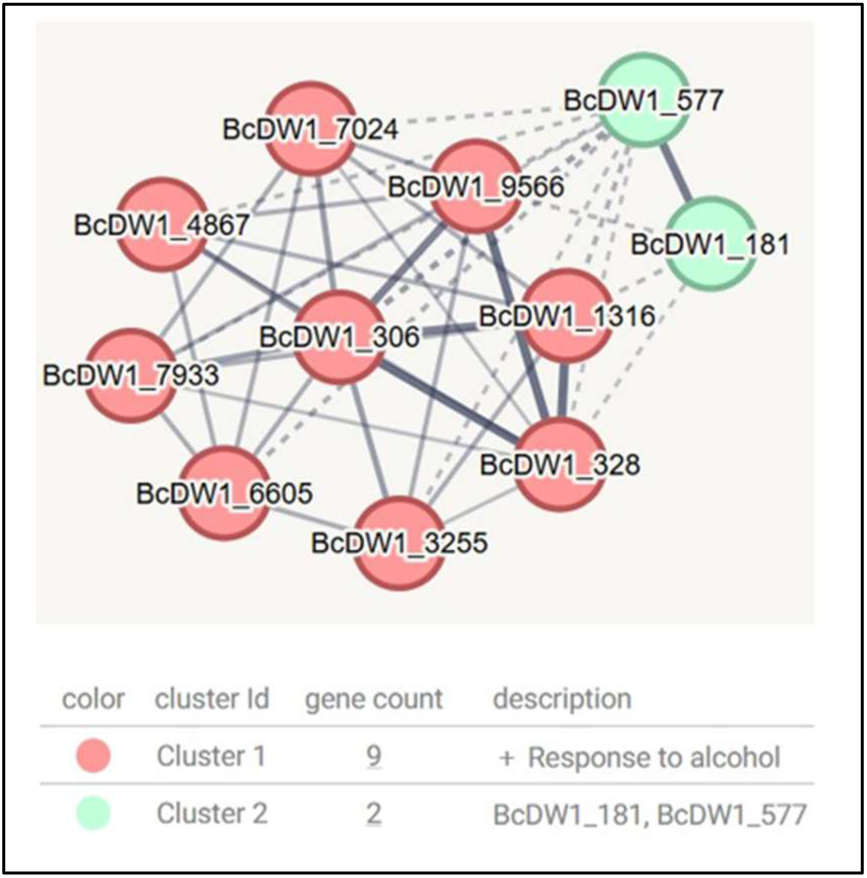
Curated PPI of the NAD(P) domain in closely related *Leotiomycetes* species (*Botrytis cinerea*) was compiled using STRING along with manual curation. Solid lines indicate experimentally verified or high-confidence predictions, while dashed lines depict functional associations derived from genomic context (such as gene neighborhood and co-expression). Protein functions were annotated utilizing resources like InterPro. This network emphasizes potential molecular complexes that play a role in alcohol metabolism. The accession number corresponding to the family of proteins is depicted in **Supplementary Table 6**.

#### 3.7.3 The activity of light-linked guanylate cyclase may influence carnitine metabolism during nutrient stress in fungal world

Currently, there is no experimental evidence directly demonstrating a physical or regulatory interaction between guanylate cyclase (GC) and carnitine acetyltransferase (CAT/CRAT) in fungi or other eukaryotic organisms. In *Neurospora crassa*, transcriptomic analyses reveal elevated expression of CRAT-1 and stress-related cyclases in response to carbon starvation (Santos-Medellín et al., 2017). In *Candida albicans*, the expression of GC (CYC1) is enhanced during morphogenesis (Biswas et al., 2007), whereas CAT expression rises during changes in lipid metabolism, both of which may be triggered by nutrient scarcity (Rodaki et al., 2009).

The hypothesis is that "Guanylate cyclase activity influences cellular energy metabolism, thereby indirectly impacting the expression or activity of carnitine acetyltransferase under conditions of nutrient stress." *Boothiomyces macroporosus* is an uncommon and insufficiently described type of fungus. It is challenging to investigate the presumed role of rhodopsin-coupled guanylate cyclase with the CRAT effector domain. We can speculate based on the literature review that Rh-GC-CRAT-induced signaling may activate stress response transcription factors that also control CRAT expression under typical conditions such as carbon starvation, oxidative stress, shifts in pH, or variations in CO₂ levels.

### 3.8 Interaction of Biosynthetic gene cluster (BGC) with photoreceptors paves unexplored way for opto-synthetic biotechnological avenues in the relevant fungal system

#### 3.8.1 Light-mediated biosynthesis of fungal terpenoids via RPEL-coupled rhodopsin

The interaction of the *Aureobasidium pullulans* BGC genes with the photoreceptors was analyzed using protein-protein interactome (PPI), developed using STRING software. The analysis of the predicted interactome shows that the terpenoid biosynthesis pathway comprising of crucial terpenoid biosynthesis such as farnesyl pyrophosphate synthase (ERG20) interact with the RPEL effector domain indirectly via protein folding complexes, transporter proteins and tRNA synthases. These interacting partners serving as a connecting link between terpenoid biosynthesis and RPEL domain (that has been found to be coupled with rhodopsin) may play a vital role in protein production, folding and transportation of proteins **(Figure 7A)**. The RPEL motif is a G-actin binding element that regulates the activity of myocardin-related transcription factors (MRTF). These transcription factors possess several functions, interacting with other transcriptional regulators such as Hippo pathway effectors (YAP, TAZ) and TGFβ-controlled Smad proteins. The MRTF along with its partner, serum response factor (SRF) after actin polymerization, directs gene expression for cytoskeletal remodelling, contractions, and several other processes (Mouilleron et al., 2012; Miranda et al., 2021). In fungi, actin is especially crucial for growth, morphogenesis and hyphal development (Berepiki et al., 2011). Although the exact functioning and mechanistic basis of the RPEL-coupled rhodopsin’s regulation of terpenoid biosynthesis remain unclear, it is hypothesized that the light-controlled RPEL may direct changes in the fungal gene expression via upregulation/downregulation of transcription factors. The changes in gene expression alters the composition and content of fungal metabolites that may play crucial role in the fungi’s pathogenicity, fungi-host interactions or motility (Galindo-Solís et al., 2022).

#### 3.8.2 Light-switchable opto-biomanufacturing of fungal sphingolipids

Similar to the terpenoid synthesis pathway, the PPI network successfully predicts the interaction of the rhodopsin with the actin-associated proteins (BTN, CLIP) regulating the function of sphingolipid biosynthetic pathway genes **(Figure 9A and B)**. The constructed interactome predicts the crosstalk of the rhodopsin-regulated BTN, ALP and CLIP proteins with sphingolipid synthesis enzymes including carnitine acetyltransferase (CRAT) that is involved in the acetyl-carnitine biosynthesis (Hynes et al., 2011). Further, other important photoreceptors such as LOV domain containing protein, photolyases and cryptochrome are also present in the developed network which may directly modulate the biosynthesis of metabolites via light. Photolyase and cryptochrome belong to the photolyase/cryptochrome family of proteins that participate in UV-induced DNA repair and blue-light dependent circadian rhythm respectively (Sancar, 2004; Kavakli et al., 2017). The predicted interactome highlights the interaction of these light-sensor molecules with the cytoskeletal proteins, ankyrin, ALP and even RPEL domain. These cytoskeletal-associated proteins are predicted to interact with several sphingolipid biosynthesis pathway proteins including carnitine acetyltransferase and sterol binding proteins. The PPI network developed in this study clearly highlights that the cytoskeletal-associated proteins play a crucial role in mediating the crosstalk between photoreceptors and terpenoid/sphingolipid biosynthesis pathway. It is hypothesized that the regulation of the actin polymerization and subsequent transcription factors by RPEL, is perhaps controlled by light. The interactome based on literature studies and experimental procedures clearly define a relationship between blue-light sensing crytochromes with biosynthesis pathway. Similarly, the existence of rhodopsin-coupled RPEL domain in fungi, as predicted in this study may influence its effect on metabolite production via cytoskeletal proteins through an unknown mechanism.

**Figure 9.**
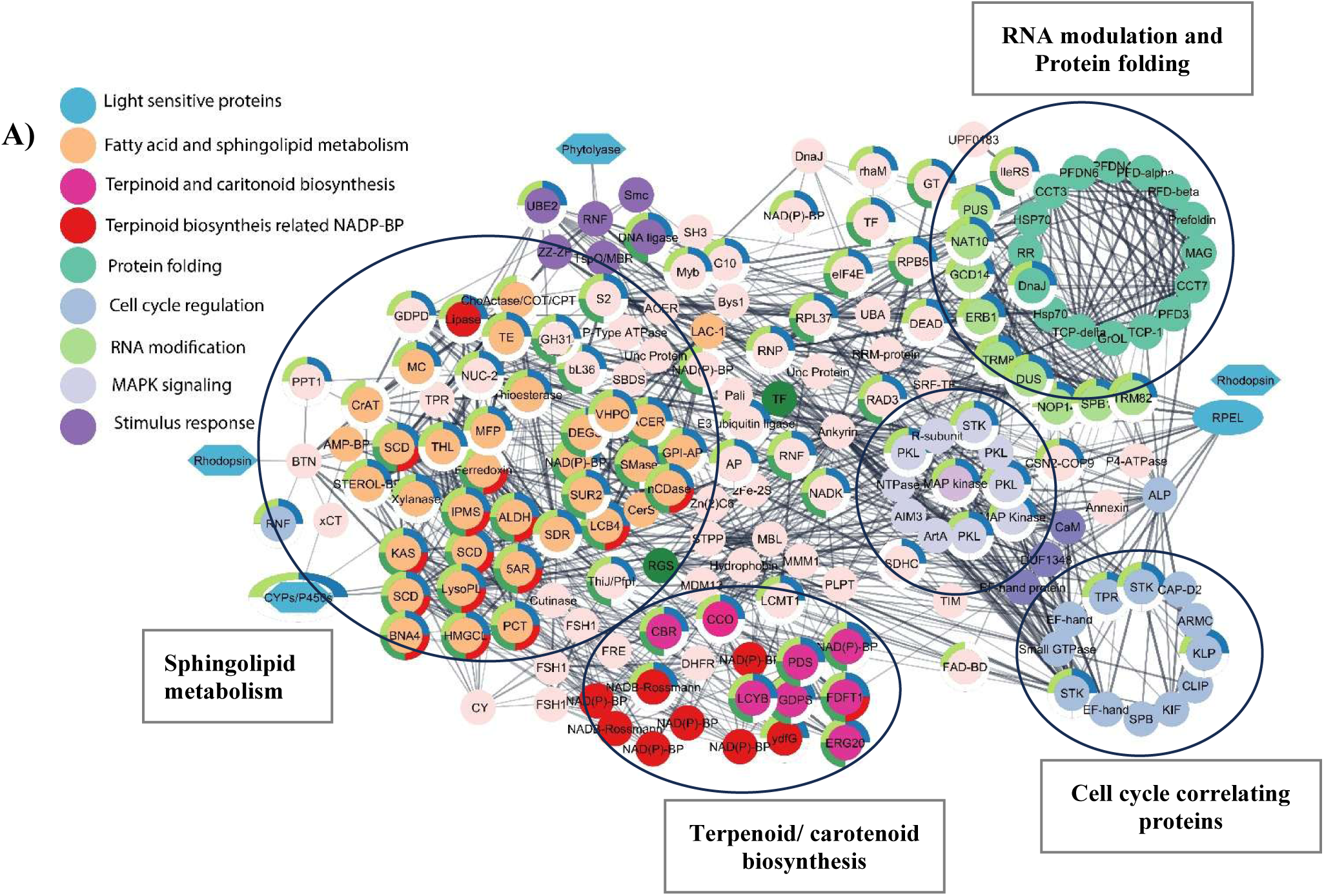
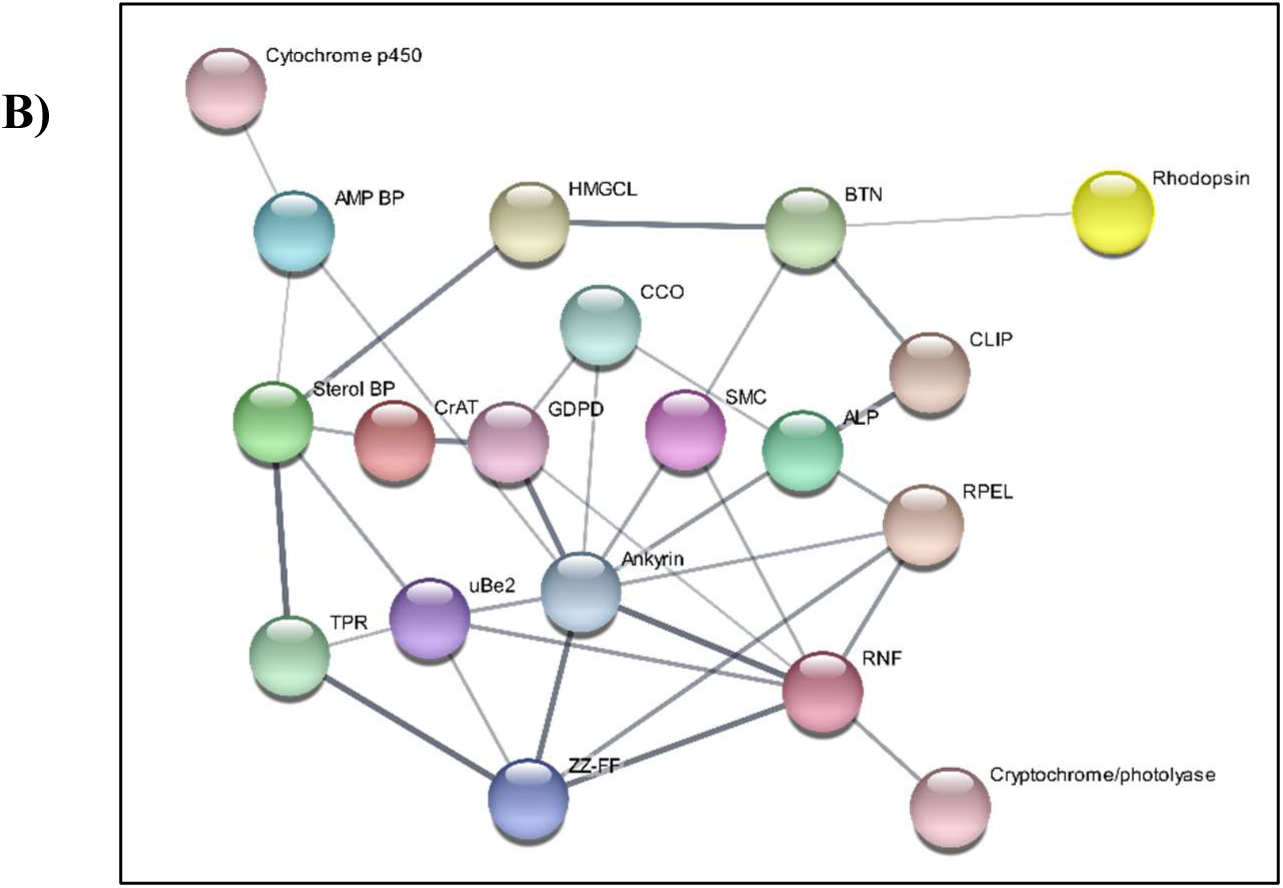
**(A)**: Opto-synthetic modulation of secondary metabolites in *Aureobasidium pullulans*. The interactome shows the interaction between light-sensitive proteins like Rhodopsin, photolyase, cryptochrome, and the biosynthesis of secondary metabolites at different signaling pathways. The important group of proteins that have been screened are related to cell cycle, sphingolipid metabolism, protein folding, terpenoid and carotenoid biosynthesis. **(B)** Manually curated protein-protein interactome showing crosstalk between light-sensing proteins and their role in the regulation of signaling cascade.

#### 3.8.3 Interaction of photoreceptor and biosynthetic gene cluster (BGC) encoding protein(s) analysis unravels novel opto-synthetic biological applications

Light being the primary source of energy plays an important role in overall metabolism in the organism across the kingdoms. Secondary metabolites have huge importance in pharmaceuticals, cosmetic, agricultural, and manufacturing sectors as antibiotics, antitumor agents, fungicides, insecticides, and perfumes (Chamkhi et al., 2021; Vaishnav and Demain, 2011). The system biology analysis conducted in this study, has depicted the interaction of direct/indirect light sensitive proteins with the secondary metabolites. The cryptochrome/photolyase (light-sensing proteins) has been seen interacting with the RPEL domain to modulate the cell cycle and the retinal/terpenoid metabolism pathways. The fungal biosynthetic cluster analysis shows the fungal system depends upon the fatty acid metabolism, such as sphingolipid and terpenoid biosynthesis pathways. The identification of novel rhodopsin-coupled RPEL domain and its plausible interaction with carotenoid and sphingolipid biosynthesis pathway highlights the role of light in fungal metabolites production. This highlights the opto-synthetic biology potential to produce commercially and therapeutically relevant metabolites from fungi by simply using light as the regulatory element. By modulating illumination conditions, the fungal metabolite content and composition can be altered that will be further used to accumulate and extract target biomolecules for human use. However, the mechanistic basis for light-based metabolite synthesis in fungi remains unexplored, that requires experimental validation to develop opto-synthetic based approaches for fungal bioactives.

## Conclusion and future perspectives

The *in silico* and systems biology analysis of the fungal genome reveals multiple novel rhodopsin-coupled effector domains. Through protein sequence alignments and phylogenetic studies, these modular rhodopsins were classified channelrhodopsin-like sequences, ion pump, or halorhodopsin. The analysis of encoding molecular domains sheds light on their role in the regulation of protein folding, transcription, DNA metabolism and gene expression. The Biosynthetic Gene Cluster (BGC) analysis of the RPEL domain coupled with rhodopsin showcases its interacting partners in the carotenoid (retinol) and sphingolipid metabolism pathway. The computational biology evidence highlights the optogenetic potential of these newly identified rhodopsin modular domains that can be harnessed for the light-mediated optobiotechnolgical and optosynthetic biological applications. Furthermore, the discovery of novel modular rhodopsin domains in fungi unearths a new branch of optogenetics that requires deep understanding and research to improve the optogenetic toolkit.

## Author Contributions

SK conceived the project, designed the outline and methodology of the analysis. ALK and ABK carried out the bioinformatics analysis and drafted the manuscript. KS and SRP generated raw data files. SM drafted and edited the manuscript.

## Author’s approval

All authors have read and approved the manuscript. The work conducted in this manuscript is original and not under consideration or published anywhere.

## Acknowledgements

The research grants (ECR/2017/000354 & CRG/2021/003158) from ANRF/SERB is duly acknowledged. Alka Kumari and Abhishek Kumar are the recipients of the Junior Research Fellowship from UGC (JRF) and ICMR (JRF), respectively.

