## Supplementary Materials for "Molecular characterization of unique multi-domain harbouring fungal rhodopsin for establishing their novel opto-synthetic biological usages"

**Supplementary Table 1:** The table provides information about the biochemical characteristics and important residues of different types of microbial rhodopsins

| **Types of rhodopsin**  (Panzer et al., 2021) | **Retinal binding site** | **Proton donor** | **Proton acceptor** | **Proton-releasing group** | **Proton pathway mediators** | **Hallmark of fungal rhodopsin** |
| --- | --- | --- | --- | --- | --- | --- |
| BR | K216 | D85 | D96 | E194 E204 D212 | R82 T89 | G116 |
| PeR | K287 | D157 | D168 | D265, E274, D282 | R154, T161 | V188 |
| LfR | K265 | D134 | D145 | D243, E253, D261 | R131, T138 | I165 |
| MdR | K258 | D127 | D138 | D236 D254, E246 | R124, T131 | I158 |
| ApR | K310 | D179 | D190 | D306, E298, D288 | R176, T183 | V210 |
| BpR | K260 | D129 | D140 | D256, E248, D238 | R126, T133 | I160 |
| EeR | K263 | D132 | D143 | D259, E251, D241 | R129, T136 | I163 |
| CapS.R | K260 | D129 | D140 | D256, E248, E238 | R126, T133 | I160 |
| TcR | K247 | D118 | E129 | D243, E235, D226 | R115, T122 | L149 |

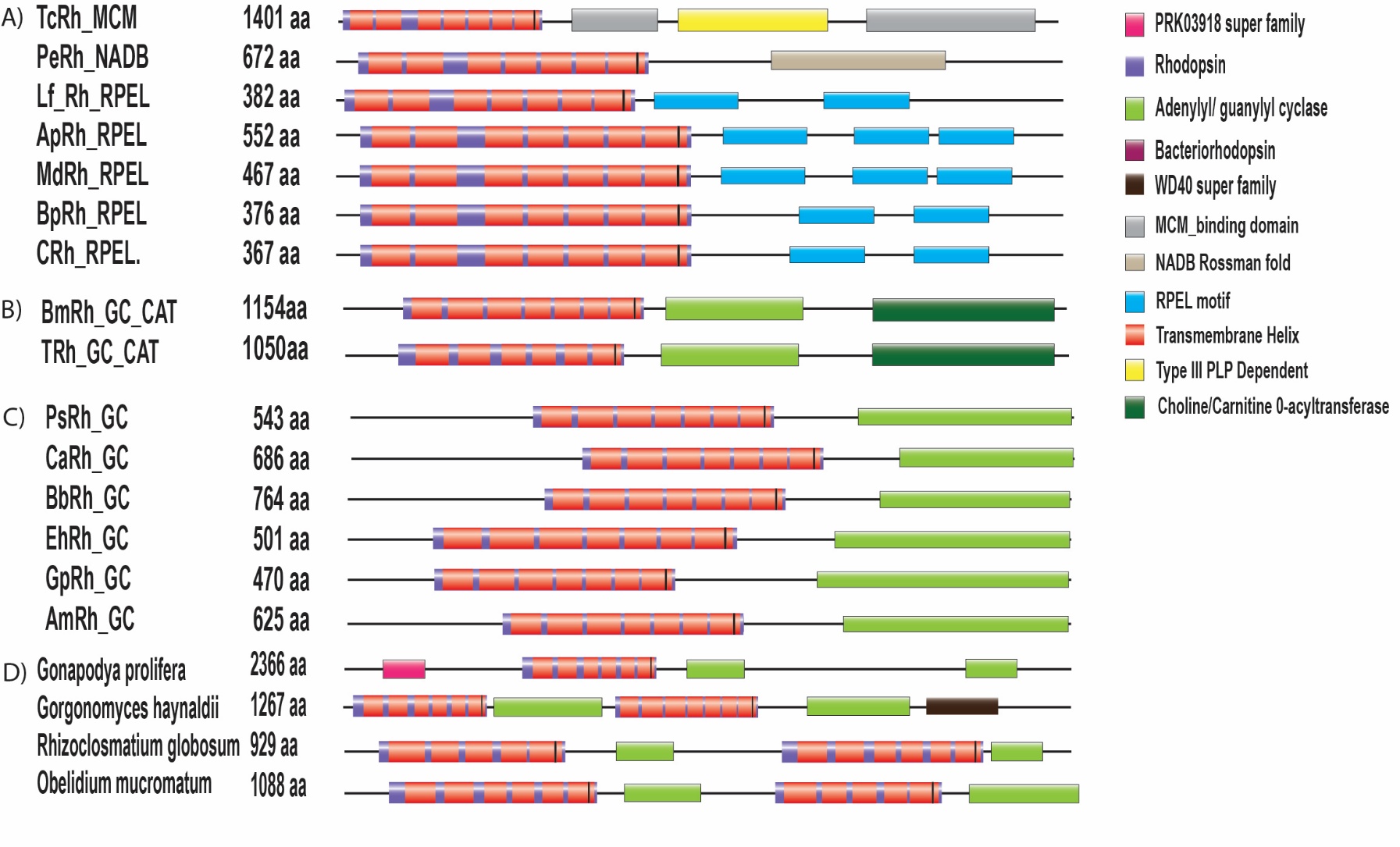

**Supplementary Figure 1:** The schematic representation of the domain organisation of twin rhodopsins (blue and orange colour) coupled with twin GC effector domains (green colour). The WD40 domain is represented by the black box. The orange zone represents the transmembrane helices, and the black line is the lysine residue. These modular domains have been found in the fungi, *Gonapodya prolifera*, *Gorgonomyces haynaldii*, *Rhizoclosmastium globosum* and *Obelidium mucromatum*.

**Supplementary Table 2:** This table details the native organisms and cellular role of different rhodopsin-coupled effector domains identified in the relevant fungal genome.

| **Domains** | **Accession No./Protein Id** | **Organisms** | **Cellular Role and Optogenetic Potential** |
| --- | --- | --- | --- |
| Rh-RPEL domain  (Modular Rhodopsin/Rh with RPxxxEL domain/motiff) | XP_007678484 | *Baudoinia Panamericana* | G-actin Binding and Sensing (Binds unpolymerized actin, senses actin dynamics), Regulation of Gene Expression (regulates nuclear shuttling and transcriptional coactivators such as MAL/MRTF-A (myocardin-related transcription factor A), which regulate serum response factor (SRF) activity (Mouilleron et al., 2008), Cell shape/motility (modulates cytoskeleton, stress fibers, and cell movement (Favot et al., 2005), Signal transduction (Links Rho GTPase signaling to actin and gene regulation (Diring et al., 2019), Protein interactions (coordinates actin and phosphatase signaling) |
|  | TIA65092.1 | *Aureobasidium pullulans* |  |
|  | KAF2681323.1 | *Lentithecium fluviatile* |  |
|  | 313850 | *Myriangium duriaei* |  |
|  | 992589 | *Capnodiales sp.* |  |
|  | KAK4972783.1 | *Elasticomyces elasticus* |  |
| Rh-MCM domain  (Modular Rhodopsin/Rh with minichromosome maintenance complex/MCM domain) | 206752 (https://mycocosm.jgi.doe.gov/cgi-bin/getDbSeq?db=Tescy1&searchTabList=protein,proteinHitDesc&hitSeqList=206752) | *Testicularia cyperi* | DNA damage response (Modulates MCM complex during replication stress/repair (Bailis & Forsburg, 2004), Genome stability (Prevents DNA re-replication, supports proper cell cycle), Cancer diagnostics/therapy (Potential biomarker and drug target in proliferative diseases) |
| Rh-NADB  (Modular Rhodopsin/Rh with Rossmann-fold^*^ domain)  ^*^The domain utilizes hydrogen bonds and van der Waals interactions to bind nicotinamide adenine dinucleotide (NAD(P)) | 2270 (https://mycocosm.jgi.doe.gov/cgi-bin/getDbSeq?db=Pseel1&searchTabList=protein,proteinHitDesc&hitSeqList=2270) | *Pseudographis elatina* | Industrial biocatalysis (Stereospecific synthesis using NAD(P)H-dependent enzymes), Drug discovery (Targets for antibiotics, anticancer agents), Protein engineering (Altering cofactor specificity for synthetic biology), Metabolic Enzymes (Rossmann fold is a core domain in many dehydrogenases and oxidoreductases, central to metabolic pathways including glycolysis, the citric acid cycle, and fatty acid synthesis). |
| Rh-GC-Carn-acyltrans  (Choline/Carnitine o-acyltransferase) | KAJ3272524.1 | *Terramyces* sp. JEL0728 | Crucial for cellular energy and lipid metabolism (Volpicella et al., 2025) |

**Supplementary Table 3:** The table showcases the effect of mutations in the retinal binding pocket of fungal rhodopsins.

| **Essential amino acid for retinal binding pocket, corresponding to BR** | **Maximum absorption shift** | **Findings** | **Impact** | **Reference** |
| --- | --- | --- | --- | --- |
| Thr89 (polar cluster near the retinal. | T89A (Minor or subtle)  T89D (Red shift)  T89V (Large blue shift) | BmRh (T89C) | Polar residue may lead to red shift. | (Marti et al., 1991) |
| Green-light-absorbing proteorhodopsins ( *λ*_max_=∼520 nm) have leucine or methionine at the position of Leu93 of BR, and leu to glutamine shifts the *λ*_max_ to ∼500 nm, | L93glutamine (blue-absorbing proteorhodopsins shift, ~500 nm), | BmRh L93M,  PeRh L93V,  ClachRh L93I | BmRh have methionine at BR93 Leu similar to green-absorbing proteorhodopsin  PeRh and have valine (non-polar) | Diversification-and-Spectral-Tuning-in-Marine-Proteorhodopsins (Man et al., 2003) |
| BR Ala215 to Thr204 in SRII from *Natronomonas pharaonis*)  And mutations in the opposite direction (serine or cysteine to alanine) result in a longer wavelength | 498 nm (sensory rhodopsin, ~70 nm blue-shifted relative to BR) | Not conclusive | ClacaRh has threonine | (Shimono et al., 2001) |
| Single nonpolar or polar amino acid at the 105th position in proteorhodopsin  (93BR) | Either green-absorbing or blue-absorbing forms of proteorhodopsin, causing a 20 nm shift toward the blue or green spectrum | PeRh (PR105 Valine)  ClacaRh(PR105 Isolucine)  BmRh, ApRh,MdRh,LfRh,EeRh, CapS.Rh,TcRh(PR105 Leucine) | PeRh(Green shift)  ClacaRh(Green shift)  ApRh,MdRh,LfRh,EeRh, CapS.Rh,TcRh(Green shift) | (Gómez-Consarnau et al., 2007) |
| Pro186 in BR is mutated to threonine in any microbial rhodopsin | Wavelength shifts to a 5–20 nm longer | Pro186 is conserved in all identified microbial rhodopsin | Not conclusive | (Inoue et al., 2019) |

**Supplementary Table 4:** Table lists the enzymes involved in the carotenoid (retinol) biosynthesis pathway in *Aureobasidium pullulans*.

| **Accession Number** | **Gene** | **Protein function** |
| --- | --- | --- |
| A0A074XGS8 | MBP1 | DNA-binding domain of Mlu1-box binding protein MBP1. |
| A0A074Y076 | PDS | Phytoene desaturase. |
| A0A074XJV1 | FDFT1 | Putative farnesyl-diphosphate farnesyltransferase. |
| A0A074XXC7 | IPPS | Uncharacterized protein. |
| A0A074YR31 | CCO | Carotenoid oxygenase. |
| A0A074XQI6 | 4HPT | 4-hydroxybenzoate polyprenyltransferase, mitochondrial |
| A0A074XG74 | GGPS1 | Geranylgeranyl diphosphate synthase 1; Belongs to the FPP/GGPP synthase family. |
| A0A074XX92 | FPPS | ERG20 farnesyl diphosphate synthase; Belongs to the FPP/GGPP synthase family. |
| A0A074XV03 | LCYB | Lycopene beta-cyclase. |
| A0A074XCF2 | DGAT2A | Diacylglycerol O-acyltransferase 2A. |

**Supplementary Table 5:** The table indicates the presence of key rate-limiting enzymes in the fungal carotenoid (retinol) pathway (Avelar et al., 2014).

| **Species** | **Phytoene desaturase (rate-limiting enzyme in the process of carotenoid synthesis)** | **Bifunctional enzyme phytoene synthase/lycopene cyclase** | **Carotenoid oxygenase** |
| --- | --- | --- | --- |
| *Baudoinia panamericana* | >XP_007674772.1 uncharacterized protein BAUCODRAFT_120791 [*Baudoinia panamericana* UAMH 10762] | >tr\|M2N1R4\|M2N1R4_BAUPA Bifunctional lycopene cyclase/phytoene synthase OS=*Baudoinia panamericana* (strain UAMH 10762) OX=717646 GN=BAUCODRAFT_66702 PE=3 SV=1 | >tr\|M2NF37\|M2NF37_BAUPA Carotenoid oxygenase OS=*Baudoinia panamericana* (strain UAMH 10762) OX=717646 GN=BAUCODRAFT_31874 PE=3 SV=1 |
| *Aureobasidium pollulans* | >XP_029765521.1 phytoene desaturase [*Aureobasidium pollulans* EXF-150] | >tr\|A0A074XV03\|A0A074XV03_AURPU Bifunctional lycopene cyclase/phytoene synthase OS=*Aureobasidium pollulans* EXF-150 | >XP_029765523.1 carotenoid oxygenase [*Aureobasidium pollulans* EXF-150] |
| *Lentithecium fluviatile* | >KAF2679432.1 phytoene dehydrogenase [*Lentithecium fluviatile* CBS 122367] | >KAF2679437.1 Lycopene beta-cyclase [*Lentithecium fluviatile* CBS 122367] | >KAF2679438.1 carotenoid oxygenase [*Lentithecium fluviatile* CBS 122367] |
| *Myriangium duriaei* | >KAF2157328.1 Phytoene dehydrogenase [*Myriangium duriaei* CBS 260.36] | >KAF2157329.1 Lycopene beta-cyclase [*Myriangium duriaei* CBS 260.36] | >KAF2157244.1 carotenoid oxygenase [*Myriangium duriaei* CBS 260.36] |
| *Capnodiales sp.* | >KAK3049059.1 ATP-binding cassette transporter CGR1 [*Extremus antarcticus*] | N. A | N. A |
| *Pseudographis elatina* | N. A | N. A | N. A |
| *Testicularia cyperi* | >PWY98742.1 putative phytoene dehydrogenase [*Testicularia cyperi*] | >PWY99893.1 Phytoene synthase [*Testicularia cyperi*] | >PWZ01013.1 LOW QUALITY PROTEIN: carotenoid oxygenase [*Testicularia cyperi*] |
| (N.A-Homologous sequence is not identified) | | | |

**Supplementary Table 6:** List of proteins of retinal synthesis and their corresponding accession number in *Botrytis cinerea* and homologous BLAST search in *Pseudographis elatana.*

| **Accession number** | ***Botrytis cinerea* BcDW1** | ***Pseudographis elatana*** |
| --- | --- | --- |
| BcDW1_9566 | Metallo-beta-lactamase family protein. | N. A |
| BcDW1_1316 | 2fe-2s iron-sulfur cluster binding domain-containing protein. | >jgi\|Pseel1\|1848\|g2010.t1 |
| BcDW1_328 | Nitrate reductase | >jgi\|Pseel1\|5594\|g5531.t1 |
| BcDW1_3255 | hort-chain dehydrogenases/reductases (SDR) family. | >jgi\|Pseel1\|7896\|g7600.t1 |
| BcDW1_6605 | Beta-ketoacyl-ACP synthases family | >jgi\|Pseel1\|7648\|g7376.t1 |
| BcDW1_7933 | Putative short-chain dehydrogenase protein. | >jgi\|Pseel1\|1534\|g1803.t1 |
| BcDW1_4867 | Putative short-chain dehydrogenase protein | >jgi\|Pseel1\|8692\|g8386.t1 |
| BcDW1_306 | Acyl-CoA desaturase | >jgi\|Pseel1\|563\|g11136.t1 |
| BcDW1_577 | Putative retinol dehydrogenase protein | >jgi\|Pseel1\|6584\|g475.t1 |
| BcDW1_181 | S-(hydroxymethyl)glutathione dehydrogenase | >jgi\|Pseel1\|1560\|g1829.t1 |
| BcDW1_7024 | Putative short-chain dehydrogenase protein | >jgi\|Pseel1\|8692\|g8386.t1 |
| (N.A-Homologous sequence is not identified) | | |

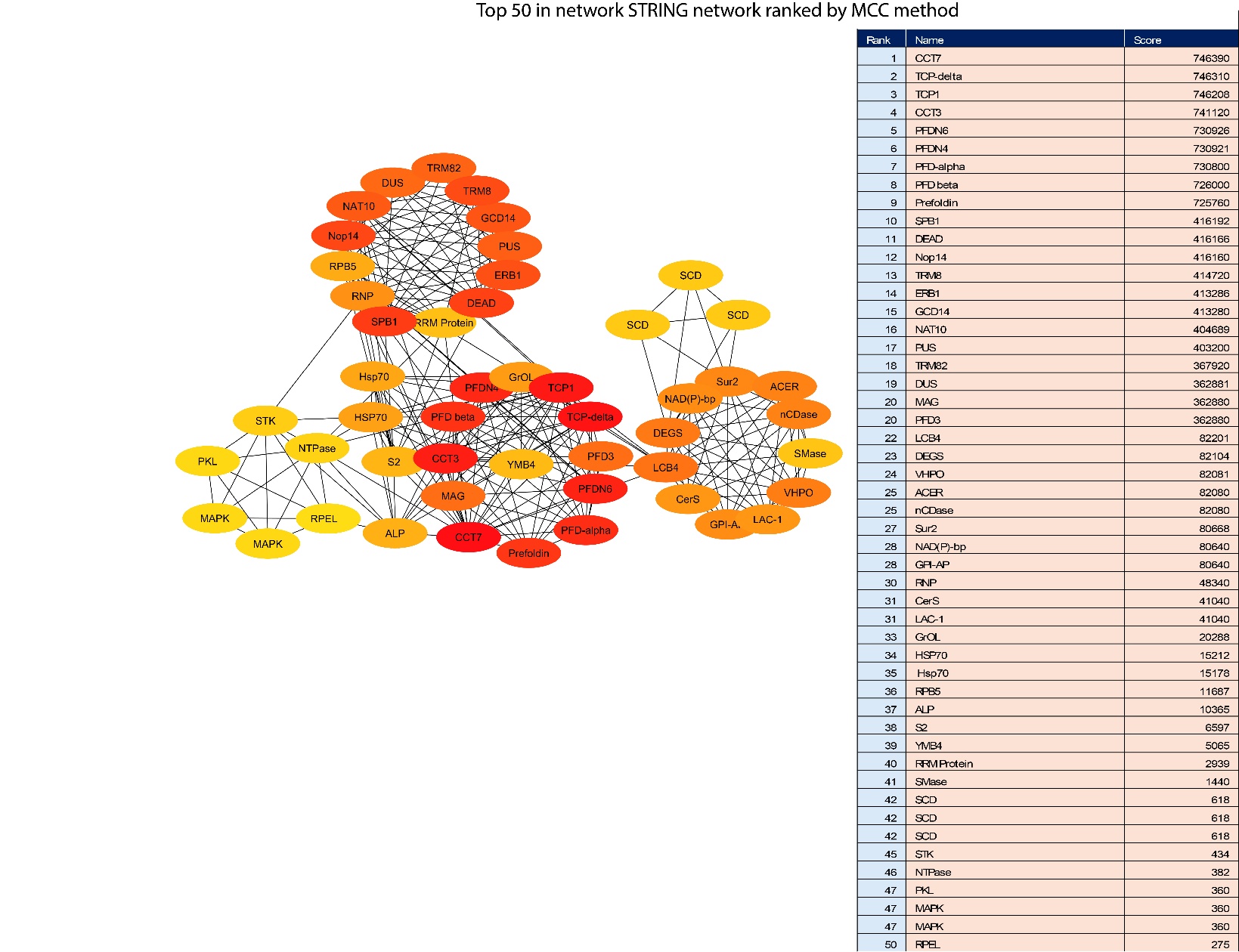

**Supplementary Figure 2:** Predicted interactome using STRING and CytoScape depicting the crosstalk between RPEL effector domain in *Aureobasidium pullulans* with protein modification and sphingolipid metabolism pathways. The principal nodes that control the overall network were analysed by CytoHubba analysis using the betweenness-centrality algorithm. Colour range (orange red to yellowish) indicates scores ranked by betweenness method.

**Protein sequences of the effector domains used in the current study:**

The following table denotes the sequences that have been employed for the study of modularity, homology, phylogenetic tree, structural and Biosynthetic Gene Cluster (BGC) analysis. The full-length sequences are also appended.

| ***Phylogenetic pattern of newly identified modular fungal rhodopsins*** | | |
| --- | --- | --- |
| **Organism** | **Domain** | **Sequence (amino acids)** |
| *Capnodiales* sp. | Rhodopsin | 39-271 |
| *Elasticomyces elasticus* | Rhodopsin | 42-274 |
| *Baudoinia panamericana* | Rhodopsin | 39-271 |
| *Aureobasidium pullulans* | Rhodopsin | 89-321 |
| *Myriangium duriaei* | Rhodopsin | 37-269 |
| *Lentithecium fluviatile* | Rhodopsin | 46-276 |
| *Pseudographis elatina* | Rhodopsin | 66-298 |
| *Terramyces* | Rhodopsin | 65-271 |
| *Cladophialophora carrionii* | Rhodopsin | 24-249 |
| *Halobacterium salinarum* | Bacteriorhodopsin | 1-231 |
| *Anabaena* | *Anabaena* Sensory Rhodopsin (ASR) | 1-261 |
| *Chlamydomonas reinhardtii* | Channelrhodopsin | 1-737 |
| *Boothiomyces macroporosus* | Rhodopsin | 65-271 |
| *Testicularia cyperi* | Rhodopsin | 29-258 |
| ***Homology analysis of the microbial rhodopsins*** | | |
| *Anabaena* | *Anabaena* Sensory Rhodopsin (ASR) | 1-261 |
| *Halobacterium salinarum* | Bacteriorhodopsin | 1-231 |
| *Pseudographis elatina* | Rhodopsin | 66-298 |
| *Lentithecium fluviatile* | Rhodopsin | 46-276 |
| *Myriangium duriaei* | Rhodopsin | 37-269 |
| *Aureobasidium pullulans* | Rhodopsin | 89-321 |
| *Baudoinia panamericana* | Rhodopsin | 39-271 |
| *Capnodiales* sp. | Rhodopsin | 39-271 |
| *Elasticomyces elasticus* | Rhodopsin | 42-274 |
| *Testicularia cyperi* | Rhodopsin | 29-258 |
| *Cladophialophora carrionii* | Rhodopsin | 24-249 |
| ***Structural analysis of microbial rhodopsin*** | | |
| *Aureobasidium pullulans* | Rhodopsin | 89-321 |
| *Anabaena* | *Anabaena* Sensory Rhodopsin (ASR) | 1-261 |
| *Baudoinia panamericana* | Rhodopsin | 39-271 |
| *Lentithecium fluviatile* | Rhodopsin | 46-276 |
| *Myriangium duriaei* | Rhodopsin | 37-269 |
| *Capnodiales* sp. | Rhodopsin | 39-271 |
| *Elasticomyces elasticus* | Rhodopsin | 42-274 |
| *Leptosphaeria maculans* | Light-gated proton rhodopsin | 49-281 |
| ***Domain analysis of the newly identified modular fungal rhodopsins*** | | |
| *Aureobasidium pullulans* | Rh-RPEL | FL: 1-552  Rh: 89-321  RPEL (1): 434-454  RPEL (2): 478-501  RPEL (3): 522-545 |
| *Baudoinia panamericana* | Rh-RPEL | FL: 1-376  Rh: 39-271  RPEL (1): 305-326  RPEL (2): 347-370 |
| *Lentithecium fluviatile* | Rh-RPEL | FL: 1-382  Rh: 46-276  RPEL (1): 308-331  RPEL (2): 352-375 |
| *Myriangium duriaei* | Rh-RPEL | FL: 1-467  Rh: 37-269  RPEL (1): 349-372  RPEL (2): 393-416  RPEL (3): 437-460 |
| *Capnodiales* sp. | Rh-RPEL | FL: 1-367  Rh: 39-271  RPEL (1): 296-316  RPEL (2): 337-360 |
| *Testicularia cyperi* | Rh-MCM | FL: 1-1401  Rh: 29-258  MCM (1): 446-565  PLPDE: 610-891  MCM (2): 902-1377 |
| *Boothiomyces macroporosus* | Rh-CRAT | FL: 1-1154  Rh: 65-271  AC: 318-492  CRAT: 516-1095 |
| *Rhizoclosmastium globosum* | Rh-TWIN | FL: 1-929  Rh (1): 116-258  GC (1): 284-465  Rh (2): 544-746  GC (2): 772-921 |
| *Gorgonomyces haynaldii* | Rh-TWIN | FL: 1-1267  Rh (1): 63-238  GC (1): 260-443  GC (2): 701-892  WD40: 902-1038  Rh (2): 473-681 |
| *Obelidium mucromatum* | Rh-TWIN | FL: 1-1088  Rh (1): 133-317  GC (1): 343-524  Rh (2): 713-856  GC (2): 907-1087 |
| *Pseudographis elatina* | Rh-NADB Rossman | FL: 1-672  Rh: 66-298  NADB Rossman: 372-639 |
| *Caternaria anguillulae* | Rh-GC | FL: 1-686  Rh: 242-454  GC: 504-684 |
| *Allomyces macrogynus* | Rh-GC | FL: 1-625  Rh: 180-397  GC: 443-623 |
| *Blastocladiella britannica* | Rh-GC | FL: 1-764  Rh: 294-506  GC: 563-734 |
| *Paraphysoderma sedebokerense* | Rh-GC | FL: 1-543  Rh: 105-311  GC: 361-541 |
| *Entophlyctis helioformis* | Rh-GC | FL: 1-501  Rh: 55-265  GC: 314-499 |
| *Globomyces pollinis-pini* | Rh-GC | FL: 1-470  Rh: 93-236  GC: 286-468 |
| ***Biosynthetic Gene Cluster (BGC) analysis*** | | |
| *Aureobasidium pullulans* | Rh-RPEL | FL: 1-552  Rh: 89-321  RPEL (1): 434-454  RPEL (2): 478-501  RPEL (3): 522-545 |
| FL: Full length sequence | | |

**The full-length sequences are as follows:**

**>****TIA65092.1 family A G protein-coupled receptor-like protein [*Aureobasidium pullulans*] (Rh-RPEL)**

MTTSSRQAGAIESSARSSSELAAALFKNVSPSAALPSHPQPLSSLHPIPSSIHAPAMIVDPVEAFKATSSVAPIPTVVPSLPEYETVTETGTRALWAVFVLMLLSMIVFVGLSWTVPISKRLYHVVTTLIVTFASLSYFAMATGHGISYHRTTVTDSHRHVPDTTHDVYRQVYWARYVDWSLTTPLLLLDLALLAGLSGGHILLAIVADVIMVLTGLFAAYGTEGTPQKWGWYAIACIAYLVVIWMLAVHGRANAMAKGGKVGKFFASIAGFTLVIWTIYPIVWGVADGSRKMSVDQEIIAYAVLDLLAKPVFGAWLLFTHQSMPETQVELGGFWTHGVSSEGAIRVDPVTTPVNDNPARFMQGLSLSGEEGAQELTHHLNSHLPAHNDICFCGSTRLPKLLYCSGTQQQAPATMSDELTPATTSPISLERRNSLEKAIQNRPEVHELREKHILLNTNAAPALQAQQQELQRHKLTDSLNKAIASRPEMDELVERNILPDSTAAPALQNHQRELAAAMRRDSIEKHLQTRPSPAELVKEGILEADENPLDGP

**>XP_007678484.1 hypothetical protein BAUCODRAFT_149869 [*****Baudoinia panamericana* UAMH 10762] (Rh-RPEL)**

MIVDPVQAFATGTIPIATSSATPLPTIIPSLPEYQTATDTGNRTLWVVFVVMLVASIVFAGLSWNVPMSKRLYHIVTCLITIFASLSYFAMATGHGIGYHHVVERESHKHVPDTTYDVYREVYWARYVDWSLTTPLLLLDLCLLAGISGGNIMIAIVADIIMILGGLFAAFGSEGTPQKWGWYTIACIAYLVVIWQLAYNGRAMAMSKGGKVGNFFAAIGGFTLVIWTVYPIIWGIADGSRNMNVDEEIIAYAVLDILAKPVFGTWLIYTHMTMPETNVEIGGFWSEGLKGEGQLRVGDDEGKQDGLAKRPEKDELVERNILPDSMAAPALQEKQRELEKHMRADSLEKHLQQRPKVEELVKEGILQPDENPIAEG

**>****KAF2681323.1 family A G protein-coupled receptor-like protein [*****Lentithecium fluviatile* CBS 122367] (Rh-RPEL)**

MIIDPVEILKKTSTAGPIPTATPTSVDPIPTVLPDSPEKQFVGDGGTRTLWVVFIVMLISSAVFAGLSWRVPVGRRLYHVITTLITIFAAISYFAMATGHGVSVHTIQVRHQIDHLPDTFTEVQRQVFWARYVDWSLTTPLLLLDLSLLAGLNGAHILMAIVADIIMILTGLFAAFGSEGTPQKWGWYAIACIAYLVVIWHLAVNGRAQAQAKGDKVGSFFLAIAGFTLIVWTAYPIVWGIADGSRNLSVDGEIIAYAVLDILAKPVFGTWLLIAHARMPETNIDLGGFWSYGLGGEGSVRLGDDDDNLKKGLQHRPDRDTLVERNILPDSNAAPALQGHQKELERHMRANSLEKGLQHRPDPETLVKKGILEEDENPLKDA

**>jgi|Myrdu1|313850|estExt_fgenesh1_pg.C_7_t10474 [*Myriangium duriaei*] (Rh-RPEL)**

MIVDPNQVAAFAAAKTSVAPVPSFTPGPSVSYETAGDSGKRALWVVFVIMLVATVVFSFLSFSVPISKRLYHVITTLIVTFAALSYFAMASGDGISLHKNVVTEEHKHVPDTQTYVYREVYWARYVDWSLTTPLLLLDLALLAGLSGGNIVIAVVADIIMVLTGLFAAFGKEESPSKWGWYAIACIAYLVIVWQLAVNGRATAFGKGGKVGTFFASIAGFTLIVWTIYPIVWGVADGARIASVDQEIIAYAVLDVLAKPVFGAWLLYTHAAIPETNIEVGGFWTHGVSGEGAIRSVAVVEFFPDLVPSLVERTHFNTLTTLMTMAANSDTTPAIGVDDSPINTLERRNSLEKHLQTRPEEQDLKNRHILLDTNAAPALQAQQHELERQRITDSLKKGLSHRPEKDELVQRNILPDSNAAPALQGKERELSKHMRANSLEKQLQNRPSPEVLIKEGILEANENPLADS

**>****jgi|Cap6580_1|992589|estExt_fgenesh1_pm.C_720012 [*Capnodiales* sp.] (Rh-RPEL)**

MLVDPVKAFATGILPLPTASASPLPSVVPSAAEYQDATETGNKTLWVVFTIMFIASIVFSGMSWNVPISKRLYHIVTTMIVIFASLSYFAMASGHGISYHHIEIRESHKHVPDTTKDIYRQVYWARYVDWSLTTPLLLLDLCLLAGLSGGHILLTMVADIIMILTGLFAAFGTEDTPQKWGWYAIACVAYLVVIWHLVIHGRATAMSKGGKVGNFFAAIGGFTLVIWTVYPIIWGIADGSRNMNVDEEIIAYAVLDVLAKPIFGAWLLFTHMSMPETDVDLGGFWSEGLKGEGNIRGLAKRPEKDDLVEKNILPDSTAAPALQERRRELERHMRADSLEKHLQARPKPEELVKEGILNANENPVAES

**>****KAK4972783.1 hypothetical protein LTR42_006077 [*****Elasticomyces elasticus*] (Rh-RPEL)**

MIDPVQAFATAMASATLPLATSSVSPIPTVVPTLPEYQDATETGTRTLWVVFVIMLLATVVFSGMAWTVPISKRLYHIVTTLIVIFASLSYFAMASGHGISYHHVVVRESHKHVPDTTTDLYRQVYWARYVDWSLTTPLLLLDLALLAGLSGGVILMIMVADIIMILTGLFAAFGTEDTPQKWGWYAIACIAYLVVIWHLVIHGRATAMNKGGKVGNFFAAIGGFTLVIWTVYPIIWGIADGSRNMNVDEEIIAYAVLDILAKPVFGAWLLFTHASMPESNVELGGFWSEGLKGEGNIRVGDDDEATKQQYEIMAEQEANNAVDRSPIAQLERRNSLEKFLQGRRPEAQELKNRHILLDTTSAPTLQARQAELERQRITDSLKRGLAKRPEKGDLVEKNILPESTAAPSLQETQKKLAKQMRADSIEKHLQGRPKQEGILHADETPIAEG

**>****KAJ3272524.1 hypothetical protein HDV01_005475** **[*Terramyces* sp. JEL0728] (Rh-GC-CRAT)**

MNQNDLYLFDRGNSDGWYKNNASAFVISGTFAFFANLTANSPDKKSLAVVLLLVNLITLCSYLLMAWRLTPQFKDGNQYPVDVARYLEWIATCPNLILLVGEFTKENTFTFRIFVYDYLMLISGFLASTTTGSISTISLLMSCIYFVLIMQGLHFMFTKAIAGGTLCVLDKQTLQIAHASTMVGWTIYIIAKAFLTIVLVNATIEQTQSETVQNISGIVATMEKEFNNTEAILEKVFPSELLPEIKEGNFPEAKDYKSVTIFFSDITNFNSITSKSSTRDVLKTLDALWTKYEEIGRKWGICKVETIGDAFLGISGCPTRTDNHAENCLNFALEIIEMAKEFKTSANDSIHIRIGIHSGPVVAGFMGTNNPRWCILGDAVNTASRIETLSKPMKINISESTFELVKEKNLFEFSEPHIYEIRGKGRFNMYWVAAKPRSINRNLVAKRFQQTFASQGKVPRLPIPTLENLEQKYLKSCQPLLAKEDYEKTEAIVKEFLSKVDHPEQPSDLLTKPPPKGVLTSFQIKRAAGLITNLLNFNDMLNNQTFPVEAIKGTPLCMNQYKNIFGTTRLPGKTSDTLSHQYPTTAKHIIVLTKNEIYKVQVLADDGKRVSNAELERVLLNIGKETLVDRHPPHIGVLTAGDRDTWYEASEKLCALSPSNVKNFDIIKDSLFAVCLDDHSTKKNLDLSLTQIFHNNNAQNRWFDKSLQLIITNSGRAGLNGEHTPSDAVVPGNVMDYIISHEPSIDPQNTVQGQYLPPPQRLEWVVDDSVNSLIEKARGVAQALIDDTESTLLQTDYYGSRFMKEIAKTSPDAYIQIALQLAYYRLHQKPTAVYESASTRFFKHGRTETGRSMSNESLEFISTFDNDDVLYDTKRELFRKAIATQSNYMKDAAYGKGIDRHMLGLRCMIKPDEMEKATMFTDPAYITSMTFRLSSSNMSPGRNFYGGFGPVVFDGYGINYAIDKDNLKFSISAKRSCTETKIYRLRDELEKVFKDLCILFPKRSEIWGKNWRKEQQAEKVADIRFAKMKTLSDEYLSKQASLAEKYLDRK

**>KAJ3254479.1 hypothetical protein HK103_007115 [*Boothiomyces macroporosus*] (Rh-GC-CRAT)**

MSNLWSAKIKNETEKSRIQSTTINQMILSLFITVPMAVYTHYVHSGAQPFDRGNADSWYNYNASIFTFTGLSAFFAMLTAKSNDKKSLAIVLGLVNLITLCSYLTIAKRLTPNVPDSNGYPVDFARYLEWIACCPNMVLLIGEFTKDAEYTSQTFLNNYVMLCCGFLASVTGESLSIIFTSITVYLFIKVMTGLHRMYSNAIDGRTGCTLDRTMVRIANITTTFGWSLFPIVWFLVRFKVISFETGEMLYCFGDIVAKAFLTLVLVNATIEHAQNETIQRISGIASVMEREMNNTDVVLEKFIPPEVLSNIKEGNFPEAQDYESVTILLSDITNYNVLSARNSTRDVLKTLDSLWSKYEEIGKKWGIYKVETIGDAYLGVSGCPTRSENHAENCLNFALDILEMTTEFKTETKEGIQIRIGIHSGPVVAGLMGANNPRWCIVGDSVSTASRIEALSKPMRINISESTYELVKDKNMFEFSGPNSYEVKGKGRSLNRNLLTKRFQQTFANQDKVPRLPIPTLENLEQKYLKSCQPLLSKEDYEKTESIVKEFLSKDGFGTTLQERLIEYEKKEPHSWLEKIWLQKAYLEWREPSMINVNWWCQLVDHPEHPLDLLTKPPPKGVLTSFQIKRAAGLITNLLNFNDMLNNQTFPAESIKGTPLCMNQYKNIFGTTRLPGKTSDSLSHQYPTTAKHIIVLTKNEIFKVQVLTDDGKRVPIAEIERVLLNIGKETLVERSPPHIGVLTAGDRDTWFEAYEKLRALSPTNVKNFDIIKDALFAVCLDDHSTKKNLDQSLTQIFHNNNAQNRWFDKSLQLIVANSGRAGLNGEHTPSDAVVPGNVMDYIISHEPSIDPENAAQGEYLAPPQRLDWVVDDSVNALIEKARGVAQALIDDTESTLLQTDYYGSRFMKEVAKTSPDAYIQIALQLAYYRLHKKPTAVYESASTRFFKHGRTETGRSMSNESLEFISTFDNDDVLYDSKRELFKKAVNTQSNYMKDAAYGKGIDRHMLGLRCMIKPDEMDKATMFTDPAYITSMTFRLSTSNMSPGRNFYGGFGPVVFDGYGINYAIDKDNLKFSISAKRSCTETKIYRFKDELEKVFKDLCILFPKRSEVWGKNWRKDQQAEKVADIRFAKMKALSDEYLTKQASLAEKYSKKA

**>jgi|Rhihy1|734731|fgenesh1_pg.8_#_202 [*Rhizoclosmastium globosum* ] (Rh-TWIN)**

MAEKDNSGAVYYVFRNFVVWSVLTVSGFYLLGTGYRGPTFDRGIADEFYAYTAASFGIAVIYSYIAYANAQNNEKRNLGKVLCIVNFIAMSSYIMQYTRTTPAFTDYVGYPVDPSRFYEWLATCPILIYLIAEITDNHHMADTTASYDYSLIILGFIACFLKQPYSEWFACGSTIFFFYTIYGLMVMFNSAIAGRTGCKLDVNSLYYARFVTFLAWNSFATVWYIQRAQIVTYEQGEVLFCVSDIFAKVFLTFILVNATLEESMNSKAKKMEAVASEIESQMAQADKLLEKLMPPSIVEAMKAGKANGAEEYASVTVFFSDITNFVALSGKSSTKDMLATLNKLWVEYDVICKRWGVYKVETIGDAFLGVVGAPDRIPDHAERAANFAIDVIRMVQDFRTVNDEEIVTRIGLNSGPITAGILGDSNPHWCIVGDAVNTASRMESTSKPMRIHISENTYKLINGKGFKLEGPDVMNIKETDWKRKIEAAKAEKDQSGTVFYVYRNYAAWMVVTLFGFYQLANGHKGPTYERGFAIEAYAYTACCFGIAVIYSFIAFFNAQNNEKRDLGKVLCVVNFISMVSYLLQWTGYTVSYTDVYGHPTDPARFYEWISTCPILIYMIAEITDNHHLADNTASADYVLLVLGYSSTFLRQPFSEYAAISAATCFFFVVKGFSSMFDRAIEGKTNCKLDAGSLWNAKWVTILAWASFPLSFFTQRSGIVTYETGEMMFCVADIWSKVFLTFILVNATLEESMNSKAKKMEAVAHEIESQMAQADKLLEKLMPASIVEAMKAGKATGSEEYSSVTVFFSDVTNFHALSNKNSTKEMLATLNKLWIEYDVICKRWGVYKVETIGDAFLGVVGAPERVPDHAERAANFAVDVISMVQDFRTDKGEELITRVGLNSGPITAGILGDSNPHWCIVGDAVNTASR

>**jgi|Gorhay1|216295|fgenesh1 [*Gorgonomyces haynaldii*] (Rh-TWIN)**

MSASKVSWKDKLDQVQHASKVSGSSISTIERNVWLWVPFTAVGAFYAAIGRKSRFDRSSADNWYYYTSSVFGIGIFFSLVALASAKSAEKRSLSQVLLWVNMIACSTYMLSATRLTYTFVDVNGYPVDVARYVEWISTCPVLILLIGEITKCGNIARTTMKYDYIMLILGFIGAITREPISFLFQMMCFAYFTLILSGLSTMYDLAIQGKTGANLDKASLITAKWATLLAWNCFTFVWFGVRYRVISFATEMESELSNSDALLQRMMPPEVIEQLKTGQAPGAEEYDSVTVFFSDITNFTVLSSQSSTKDMIGTLNKLWLEYDAIAKKWGVYKVETIGDAYLGVVGCPTRTPDHAAAAVNFALDIVEMVRGFKTEMGSSIQIRVGLNSGPITAGVLGDLNPHWCIVGDTVNTASRMESTSKPMHIHISESTYKLATKFGKFKITGPDVLQVKALPHRQKALRFDRGDGPFYDMWVASIFAFGALFALLAQGSARTAEKVALTKVLFMCELLSMSTYLIQAYRISVSLPSWNGWPVDVARFLEWITTCPILIQLISDVCKTPDLTHKTVTFDYILLVCGFMASITRNPYSFLFSWIAFGCFTQVVTGLWSMFQRAIDGNTACKLHPSALRQAQLATCVSWACFPTVWLLMQFKVVSYGTGELLYGIADIFAKVVLTLILVNATVEQAQSERVSALESIATDMEKELGNTGALLSRMMPPEVIEQLKSGRAPGAEEYDSVTVFFSDITNFTVLSSQTSTKDMLATLNKLWLEYDAIAKKWGVYKVETIGDAYLGVVGCPTRSPDHAANACSFALDIVEMVRGFKTEMGSSIQIRVGLNSGPITAGVLGDLNPHWCIVGDTVNTASRMESTSKPMHVHCSESTYKLANPSGRFRFAGPDVLQIKRGLLDAVNCLRGGHLGTVSSLVVVFESGLVDVYDLSEAQERLILSFDNRVSTWGIAIQSERQWIAVSSNSHFITLYDLKTGTQRQFQGHDHNIPGIEFMGDYLVSCSIDGTCRIWNGSDGQLLHTASHGEEWKWCVRHLSKTDAQVVDKHRIQRLKDDPIDVPLLERFQSDPVLQSDCETGSGFGSPDQRSVDSMESVQHSPLKRKSDLEEQIGSPKKQELSKIEEPPIPQLPTLRSLLEQDIPEDLEEMDEDWSQQNLSETESLEYESSIEEWVTLRPKGEISTNDLILLTSKNHVYLLDQELRLICGLSHHAQWDIRYGLERNLFDTRLIYGRVDSQEASASDSKSERPPDVCQARSAWVQHRS

**>jgi|Obemuc1|938368|gm1.9661_g [*Obelidium mucromatum*] (Rh-TWIN)**

MDDIQRGNCDNCNCSVYQVQTGTRRCRGCNHGAIFHQVVVRVAPKTSGKTNWKDKLQAAMAEKDNSGAVYYVFRNFLVWSGLTGIGWYLLTTGYVAPRFDRGYADEFYAYTAACFGIAVIYSYIAYANAQNNEKRDLGKVLCIVNFIAMSSYLMQWTRTTPTFSDYVGYPVDPARFFEWLATCPILIYLIAEITDNHHMADLTASYDYTLIVLGFIACFLKQPYSEWMACCSTIFFYYTITGLMEMYSKAIDGKTGCKLDVASLKVAKVITFLAWNSFTITWYIQRAQIVTYEQGELIFCINDIFAKVFLTFILVNATLEESMNSKAKKMEAVAQEIENQMAQADKLLEKLMPPSIVEAMKAGKASGAEEYSSVTVFFSDITNFVQLSSNNTTKDMLANLNKLWVEYDVICKRWGMYKVETIGDAFLGVIGAPDRIPDHAERAANFAIDVIRMVQDFRTIKDEEIVTRIGLNSGPITAGILGDSNPHWCIVGDAVNTASRMESTSKPMKIHISENTYKLINGKGFKLEGPDVMNIKEEKAMDEIQRGTCDNCNCSVFQVQTGTKRCRGCHHGAIFHQVIVRAVPKSGKTNWKEKLEAAKAEKDNSGAVYYVFRNFIAWSVMTGLGWYLLATGYALSAFDRGYADEFYAYTAGCFAIAVIYSYIAYANAQNNEKRDLGKVLCIVNFIAMATYILQFTRTSPSFKDYVGYPVDPARFFEWIATCPILIYLIAEITDNHQMADQTATYDYVLITFGFFACFLKQPYSEWMACSAVIFFFYTIIGIVTMYTRAIEGKTSCKLDVPSLAAARAITFLAWSAFPVTWHLQRAQIVTYEQGEVMFCVSDIFAKVFLTFILVNATLEESMNSKAKKMESVAHEIEAQMAQADKLLEKLMPPSIVEAMKAGKASGAEEYSSVTVFFSDITNFVALSNKNSTKDMLANLNKLWIEYDVICKRWGMYKVETIGDAFLGVIGAPDRIPDHAERAANFAIDVIRMVQDFRTIKDEEIVTRIGLNSGPITAGILGDSNPHWCIVGDAVNTASRMESTSKPMRIHISENTYKLINGKGFKIEGPDVMNIKGKGTMNTYWLNGR

**>****jgi|Pseel1|2270|g191.t1 [*Pseudographis elatina*] (Rh-NADB Rossman)**

MRDPMEALMTMAVKTSSSSLLVPTSSLSIPTSSMTSSLLIPTSTPTSVAPIPTVKPDLPVYETVADAGTTTLWIVFVIMLFSTVAFATIAYKTPVQKRLFPVLTTFITLFATLSYFAMATGDGNSFTHIILRETHKGDIPTTVKHVFRQVFWARYVDWSVTTPLVLLDLSLLAGLNGANIIVAVVADVFMVLTGLFAAFGHSDGQKWGYYAMACIAYLSIIYLLAVPGRKAVSAKSKPTRTLFVSIALYTLVLWTLYPIIWGFGDGSRILSVDSEIIAYAVLDVLAKPGFGIWLLVAHSRIGSTTIEGFWAHGLAAEGIIRIDDDEGEGGWLVEGLQALEVVGPTRPFRIDFVKSQFKKLPYPTADFESQCIIVTGANTGLGREAARHFVRLGAKQVILGVRNLDKGKAAQQDIEATTKRTGVVEVWEVDLTSYESVKAFCARTDKLPRLDIVVENAAVAVPFFELAEGNELTMTVNVISTFLMALLLLPVLRRSSVQFNTTPRLTIVASDAHELAKFPEATAPAIFPALSDPTAKNQDDRYPTSKLIEILMVRELGPLINGSGEGKRQIILNCLTPGLCYSDLSRHATFPFSLVVAIGKTLLGRDTEVGSRTLVSAAVAGEESHGQYMADCVVYHPSKWVTGEKGAKAQKKVYAELMSILEGIYPGIAKNI

**>Q8RUT8_ChR2 [*Chlyamydomonas reinhardtii*] Channelrhodopsin**

MDYGGALSAVGRELLFVTNPVVVNGSVLVPEDQCYCAGWIESRGTNGAQTASNVLQWLAAGFSILLLMFYAYQTWKSTCGWEEIYVCAIEMVKVILEFFFEFKNPSMLYLATGHRVQWLRYAEWLLTCPVILIHLSNLTGLSNDYSRRTMGLLVSDIGTIVWGATSAMATGYVKVIFFCLGLCYGANTFFHAAKAYIEGYHTVPKGRCRQVVTGMAWLFFVSWGMFPILFILGPEGFGVLSVYGSTVGHTIIDLMSKNCWGLLGHYLRVLIHEHILIHGDIRKTTKLNIGGTEIEVETLVEDEAEAGAVNKGTGKYASRESFLVMRDKMKEKGIDVRASLDNSKEVEQEQAARAAMMMMNGNGMGMGMGMNGMNGMGGMNGMAGGAKPGLELTPQLQPGRVILAVPDISMVDFFREQFAQLSVTYELVPALGADNTLALVTQAQNLGGVDFVLIHPEFLRDRSSTSILSRLRGAGQRVAAFGWAQLGPMRDLIESANLDGWLEGPSFGQGILPAHIVALVAKMQQMRKMQQMQQIGMMTGGMNGMGGGMGGGMNGMGGGNGMNNMGNGMGGGMGNGMGGNGMNGMGGGNGMNNMGGNGMAGNGMGGGMGGNGMGGSMNGMSSGVVANVTPSAAGGMGGMMNGGMAAPQSPGMNGGRLGTNPLFNAAPSPLSSQLGAEAGMGSMGGMGGMSGMGGMGGMGGMGGAGAATTQAAGGNAEAEMLQNLMNEINRLKRELGE

**> 1XIO_*Anabaena* Sensory Rhodopsin [*Anabaena*]**

MNLESLLHWIYVAGMTIGALHFWSLSRNPRGVPQYEYLVAMFIPIWSGLAYMAMAIDQGKVEAAGQIAHYARYIDWMVTTPLLLLSLSWTAMQFIKKDWTLIGFLMSTQIVVITSGLIADLSERDWVRYLWYICGVCAFLIILWGIWNPLRAKTRTQSSELANLYDKLVTYFTVLWIGYPIVWIIGPSGFGWINQTIDTFLFCLLPFFSKVGFSFLDLHGLRNLNDSRQTTGDRFAENTLQFVENITLFANSRRQQSRRRV

**>1KGB_1 BR Bacteriorhodopsin [*Halobacterium salinarum*]**

QAQITGRPEWIWLALGTALMGLGTLYFLVKGMGVSDPDAKKFYAITTLVPAIAFTMYLSMLLGYGLTMVPFGGEQNPIYWARYADWLFTTPLLLLDLALLVDADQGTILALVGADGIMIGTGLVGALTKVYSYRFVWWAISTAAMLYILYVLFFGFTSKAESMRPEVASTFKVLRNVTVVLWSAYPVVWLIGSEGAGIVPLNIETLLFMVLDVSAKVGFGLILLRSRAIFG

**>jgi|Catan2|1255425|CE255425_1643 [*Caternaria anguillulae*] (Rh-GC)**

NTCPWISHSNSWESRPTAKRILLQPTYRPACCISPPSPTHRPHRSHHSITAHTHTHTHDGMKDKDNNLRGACSGCSCPEYCYSPTSTLCDDCKCSVTKHPIVEQPLTRNGSFRSSGASLLPSPSQPNIKVTGSSTASSNANMRNRQNNSLSVSNVRSTSSASSSNVSSPANSRPGSPSKQSALQQYQTNIADMWSWDMMLSTPSLKFLTGQFIMWAILTVAGAFYALFIQERQAYNRGWADIWYGYGAFGFGIGIAFSYMGFAGARNPEKKALSLCLLGVNIIAFSSYILIMLRLTPTIEGTLSNPVEPARYLEWIATCPVLILLISEITQADHNAWGVVFSDYALVVCGFFGAVLPPYPWGNLFNILSCAFFSFVVYSLWRSFTGAINGETPCNIEVNGLRWTRFSTVTTWTLFPLSWFAFTSGMLSFTMTEASFTMIDIGAKVFLTLVLVNSTVEQAQNQKVEAITAIAEELESQITNCDAILQKMMPEGVLEQLKNGQATEAKEYESVTVFFSDITNFTVISSRTSTKDMMATLNKLWLEYDAIAKRWGVYKVETIGDAYLGVTGAPEVVPDHADRAVNFALDIIEMIKTFKTATGESINIRIGLNSGPVTAGVLGDLNPHWCLVGDTVNTASRMESTSKAGHIHISDSTYQMIKGKFVTQPLDLMEVKGKGKMQTYWVTARK

**>jgi|Allma1|3052|AMAG_07932T0 [*Allomyces macrogynus*] (Rh-GC)**

MKDKDNNLRGACTACTCPEYCFSPSSTLCDDCKCPITKHPVVEPLSRNGSFRSSGASLLPSPSAVNVLKVGGGSAGSSVLRNRKDGPVKSSSSMLGGSRSSSPNKARASSPNGGDNDTKMTMDEFRANLQEMASWEMMMSTPSLKFLTVQFAVWLTVTVLLALYTVVAHERPKFNRGWADIWYGYGAFGFGVGVAYAYMGFTSAKSPEKRALSLCLFGVNLISFSSYVLILLRLTPSLVGTFGNPVEPARYLEWMGTCPVLILLISEITRFPHDPFKVVFHDYFLNVMGFFGAIMPPQPWGDLANILSCLGFSYVVYSLWMCFTGAIDGDTDTSVAKSGLQWIRLSTLVTWTMFPVVWFSYTTQLISFTMTEAGFVLTDIGAKVFLTMVLVNSTVEQAQNDKVEAITAIAEELEQQMTNSDAILQKMMPADVLEQIKSGQATEAQEYESVTVFFSDITNFTVISSRTSTKDMMKTLNMLWLEYDAIAKKWGIYKVETIGDAYLGVAGAPDRVPDHAERCVNFALDILDMIRAFKSATGESINIRVGLHTGPVTAGVLGDLNPHWCLVGDTVNTASRMESTSKAGHIHISEDTYKKIKDKFVTQPLDVMDVKGKGKMQTYWVLGRK

**>jgi|Blabri1|435766|estExt_fgenesh1_pg.C_10028 [*Blastocladiella britannica*] (Rh-GC)**

MSTQAIIEPAIATLLSRQQRVLLLPNPGSPTVILTPPSPRRSRQNSLDSSTPTTDLAATVAEVTPQEPQSLDEGVYQTPTSPPPLVHTEDFSAPCEVGVRSEPIVPLASSPTTPTTSTTTTTTAIMKDKDNNLRGACGSCNCPEYCYQATSSTCEDCKCAATKHPIVEQPLTRNGSYRSSGGSLLPSTSVSSASVKITPSASNIRSRVNSAMLKSVPSAASDINMTGKGKPSSTLSQYQADLGEQWSWSLMMSTPSLKFLTVQFTIWSVVTVVLMLYTLFAHEKQAYNRGWADIWYGYGAFGFAIGISFAYMGFQSARNPEKKALSLCLLGVNTISFCSYLLIMLRLTPTLEGVLSNPVEPARYLEWIATCPVLILLISEVTQFPHDPFKVIFMDYSLVLAGFVGAVIPMQPYGNLFELISCAAFSYVVYSLWIAFTGAINGETQSNVEKSGLRWIRASTVLTWSMFPVTWFAFSSGVISFTVAEAAFSMIDIGAKVFLTLVLVNSTVEQAQNMKVDVITAIAEELESQVNNCDAILQKMMPEGILEQLKNGQATEAKEYDCVTVFFSDITNFTVISSRTSTKDMMATLNKLWQEYDAIAKRWGVYKVETIGDAYLGVTGAPEESPDHAERAANFAIDIIDMIKSFKTSTGESINIRVGLNSGPVTAGVLGDLNPHWCLVGDTVNTASRMESTSKAGHIHISESTHNLIKGRFTMQPLDIMEVKGKGKMQTYWLLGRKLSVFETTFPLWESNVNEHVQMAEDPS

**>jgi|Parsed1|256095|CE256094_3907 [*Paraphysoderma sedebokerense*] (Rh-GC)**

MPAETKALKDSNVSSTAVATSKSTSLRKRGLKSAENEGGSKSLSSIGSSYLGFNGEGSGSLKFFITNFVSWCIITISITIYTIFFHEKQEYNRGNADVFYGYAAFGFFIGMSFAYMGYSGARNPEKKALSLCLLGVNFIPFVSYILLLLRLTPTIEGPVLSQPIEPARYLEWISTCPVLIFLISEVSRSKHDPVKVVLNDYLLVVTGFLGAVLPPRPWGDFFNLLSCFHFAYVVMSLWSFYTAGINGETDTSVEKGNLKVLRFSTILTWTLFPVSYFSYTSELLSFTVAEAVMCTADIGAKVFLTLILVNSTVEQAQNERVDEITAIAEELEEQIGNADKILQKMMPENVLEQIKNGGSTEAEEYESVTVFFSDITNFTVISGRTSTKDMMKTLNLLWQQYDAIAKKWGLYKVETIGDAYLGVVGAPNRCYDHAEKAVNFALDILEMIKTFKTVTGESINIRIGLHSGPVTAGILGDLNPHWCLVGDTVNTASRMESTSKAGHIHISDSTYQFIKNKFVCEALELMEVKGKGKMQTYWVFGRK

**>jgi|Enthel1|248197|CE248196_20510 [*Entophlyctis helioformis*] (Rh-GC)**

MAQATGGNKFEKSQALESMKRNFIGWTAITLPLVAYSLTIQTRLKKFDRGSADTWYLYAASAFFLGAAFAWLARLTARSNEKKSITSALLLVDMVPIATYLMQAFRLTPALRDSNGYPVDTARYLEWISTCPVLILLIGEVTKSPTVCRRTVLADYIMLVTGFLASVTREPFSSLFGTVSCACFFTVISGLWEMFTGAIEGLTASKLDKITLQAARTASIYAWSFFPIVWYSVKYKLVSFGTGEIGYCLADIVAKVFLTLVLVNSTVEQVQNERVDALSDIANALEQELSSTDKLLQRMMPAEVIDQIKSGRATEAQEYESVTVFFSDITNFTVLSSQTSTKDMLATLNALWIEYDAIAKRWGVYKVETIGDAYLGVVGCPERVPDHASRAINFAIDIMAMIRNFRTAMGSPIQIRIGLNTGPITAGVLGELNPHWCIVGDTVNTASRMESTSKAMMIHISESTYEAGKKYSPGVFDVSDPDVMQVKGKGSMVTYWVHGRK

**>jgi|Glopol1|131237|CE131236_19322 [*Globomyces pollinis-pini*] (Rh-GC)**

MIIGVLAFYAVTKRQPKYDREESNMWYFYTASVFAIGVFFACIAYFSARTKEKKSLAMVLFWCDFIAMSTYLLSATRMTHALRDINGYPVEVARYVEWLSTCPVLILLIGEITKCPEIAAETMEYDYIMLILGFIGAVTREPISFFFQMVCMAYFALVIFGLNKMFTKAINGETGCTLDPNSLKSAKIFTLLSWNAFALNWFCVRYEIYSFATGEMLFGLSDICAKVLLTLILVNATVESAQNERVNALSTIASDMETELNNSDALLSKMMPQEILDQIKAGKATEAQEYDCVTIFFSDITNFTVISSQTSTKEMLATLNALWVEYDKIAKKWGMYKVETIGDAYLGVTGCPERVPDHAIRATQFSVDIMAMISQFKTAMGSSIAIRIGLNSGPITAGILGTENPHWCVVGDTVNTASRMESTSKPMMIHISESTQKLIAKSGIFNISEPDVLNIKGKGSMVTYWVHGRL

>**XP_008722421.1 uncharacterized protein G647_00796 [*Cladophialophora carrionii* CBS 160.54] Halorhodopsin**

MGNDVFRHNGFTNTVTTNNHITAPGSDWYWTVCAVMTVSAFGFMIHSYFKPRSQRLFHYLNATICLIAAIAYFCMGSNLGWTAIEVEWVRSSPEVRGNMRQIFYVRYINWFITTPLIVVQLLLVAGLPTPTILYTLLMTEIVVINGLVGALVKSSYKWGFFTFGAVAFFFVAFAIVWDGRAYARVLGADVMRIFNILAAWIILLWTVYPVIWGVSEGGNIIPPDSEAVSYGVLDLLTKPVFGAVLIWGLRNVDLERLGIHVNDANPRVPRAAPAPKTEKDAEAAAAANNGVTAPAAATTPETAV

**>jgi|Tescy1|206752|fgenesh1_pg.7_#_159 [*Testicularia cyperi*] (Rh-MCM)**

MEAISDFAKRAGNEALSVNRPVADIDITTAGSSFLWAVFSVMAATGLGTMVWSLKVSRGERAFHYLSAAILATASVAYFAMASDLGATPVLVEFYNYAGDAAGGARPTRSIWYARYIDWTITTPLLLLEILLVSGLPLSTVFITIFFDLVMIITGLIGALVESTYKWGFYTFGCVAMFYVFYILYVPGLKSASHLGDDFKKAYLYSAMILTGLWFLYPIAWGLADGGNVISPNGEMVFYGVLDLLAKPGFALFHLFSLRRCNYSSLHLKSGKFSDYEDLGAAHYRNMRDGKAAEAGLAGDHHTTNVNGTTMGTGSTIEPAPAMRQAQVTHSLFRPQPCLCTTEKLSRGYLFPPYSSRSYILTIWHWIVPEMRTGGGMIRSIGLALATSPKRIIRRSFATSLRTARMVPDTEFTSALAKPLETVQSIYAKAQDKSAASVVKVVESHFRDTFSSDDARAKIPTLSPSSIASLLAPPSSNTAPSASHPLVRFRCMVQDTGLGTEVFLASHTQDGVEHTGLFGGEARFPVPSADESNGQQAFSNDNLTERTIMYAVSTPGQTEWAARAHRDRSGGPSHSSRSPASSSSTSSDGDISAQLANLSLMEDPPAPQKVQEVRERIRRAVDKGKSLGSTGKEPRLVAISKLHPPSAILAAHRKAGQLHFGENYVQEMVDKAKVLPREIRWHFVGGLQSNKGKLLASIPNLYLLETLDSIKAANVLQKALSSPDAAKRDEPLQVYLQVNTSGEDAKSGLPPITSADDDGKQSALLDLAVHVITKCPNLRFRGVMTIGAATNSANVQGEALEPKSVDDVVKANPDFERLIQTRRNLVRLLRSDDRIKASNESQVKEAYTELLDGSDSSADGGLELSMGMSADVDVAIMAGSDNVRVGTDCFGRRPGTRDEAMTGMKRELEIGPEQALVELRNQVQGAQQSTTSSNSAAIHDDGQPTQTASSRSAYPQKSPVPESNHIGALVKFNDLEAAESFKTAELIDVVGILDTGSLPQIEWQDTGAGQQGSSSEAPQVPCVHAILANSVDLNDVPAAWQAGSASFTSALSSKETRAELIEYIAGALAGDNLAAELVLLSITARIHARRAGLCLGALSLNISNFPAPPSSQASSSSSSSSSGGSVPETELYRRLAQLLPALVDIPMDLGSLNDPSKSLFPRSSGEGVGLEAGRLQLPSGTTIVINEGGMREGQLQDAGIRNIRALSFVLESHKLPYAFPYSEFEFDTDLNAVILSQGKSFLPFDIQCPLQAQSSDVQNGLYSPSSSAQSTAVPEDKLAQWRRSLLEARSLKTAQTFQIPESVSEHIQKEFVEDRRRQQAASSAAVAESHGGGGKSDAASGQEDLLRRMALVRLLAISRAQSSLTVETWNAAVELDTRLNERIQAQNQNQQSAASQPSISR
